# Cellular and molecular events organizing the assembly of tertiary lymphoid structures in glioblastoma

**DOI:** 10.1101/2024.07.04.601824

**Authors:** Alessandra Vaccaro, Fan Yang, Tiarne van de Walle, Saskia Franke, Roberta Lugano, Anna Dénes, Anastasia Magoulopoulou, Inés Cid-Fariña, Anja Smits, Lene Uhrbom, Sylwia Libard, Francesco Latini, Magnus Essand, Mats Nilsson, Liqun He, Thomas Olsson Bontell, Asgeir S Jakola, Mohanraj Ramachandran, Anna Dimberg

**Author notes:** Corresponding authors, &.

## Abstract

Tertiary lymphoid structures (TLS) are ectopic lymphoid aggregates associated with improved prognosis in numerous tumors. To date, their prognostic value in central nervous system cancers and the events underlying their formation remain unclear. Here, we find that TLS correlate with improved survival in glioblastoma patients. Furthermore, combining spatial transcriptomics of human tissues with longitudinal studies in murine glioma, we establish that the assembly of T cell-rich clusters is a prerequisite for the development of canonical TLS, and show that CD4 T cells play a central role in this process. Indeed, we provide evidence that IL7R^+^CCR7^+^ Th1 lymphocytes with lymphoid tissue-inducing potential are recruited to TLS nucleation sites, where they can initiate TLS assembly by expressing lymphotoxin β. Our work defines TLS as a potential prognostic factor in glioblastoma and identifies cellular and molecular mechanisms of TLS formation. These findings have broader implications for the development of TLS-inducing therapies for cancer.

**Graphical abstract:** 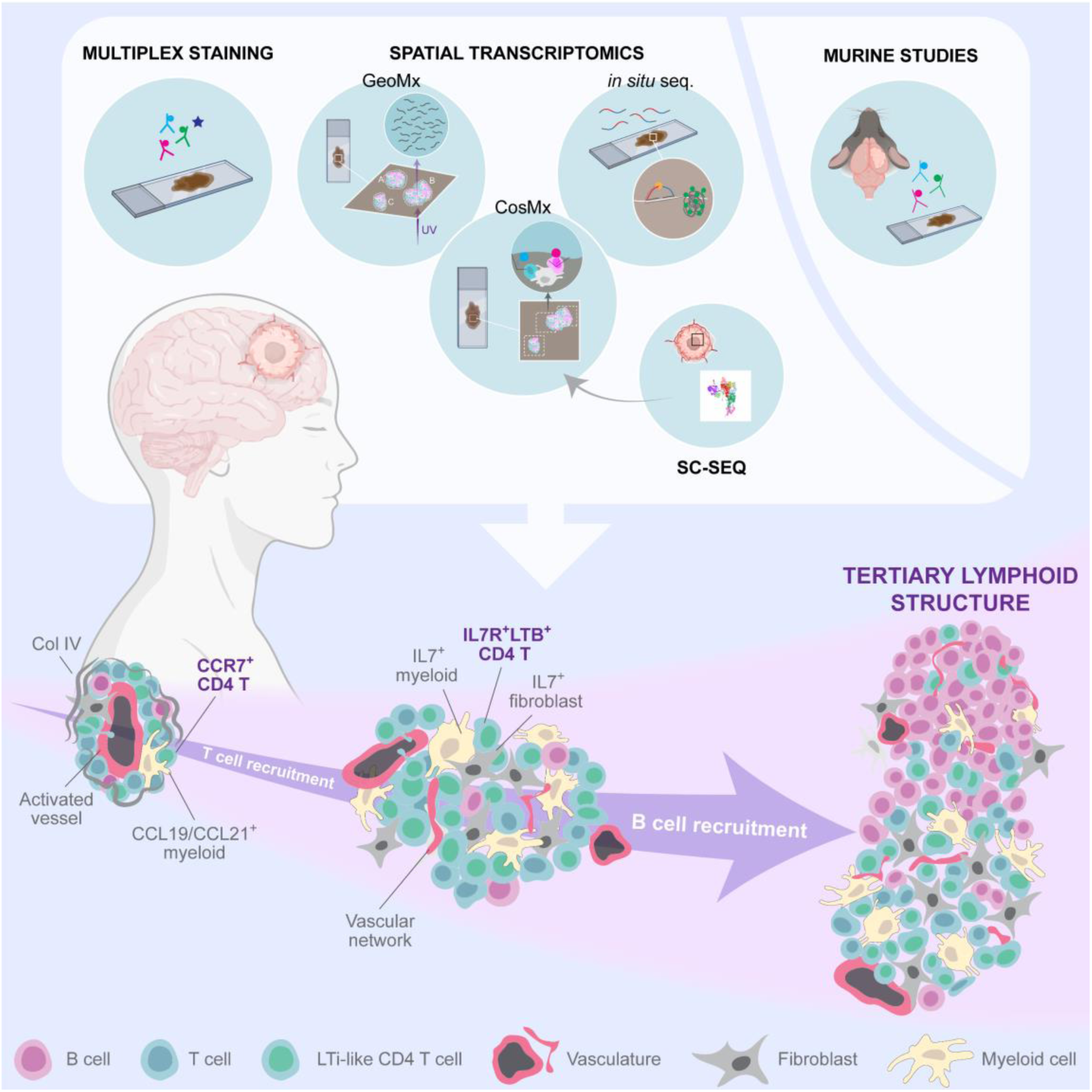

## Introduction

Tertiary lymphoid structures (TLS) are ectopic aggregates of lymphoid cells that form in pathological conditions including infections, autoimmune disorders and cancer. Due to their resemblance to the lymph nodes in terms of composition and organization, TLS are believed to be local environments where anti-tumor T cells can potentially undergo priming and activation^1^. In tumors developing outside the central nervous system (CNS), the presence of TLS has been correlated with improved prognosis and increased T cell infiltration^2,3^. These structures can display different levels of maturity, starting from immature aggregates of B cells and T cells, to primary follicle-like structures that include follicular dendritic cells (fDCs) but no germinal centers (GCs), to secondary follicle-like structures positive for both fDCs and GCs^4,5^. Interestingly, TLS maturity has been positively correlated with improved prognosis in cancer patients^4^, suggesting that boosting TLS formation and maturation could be a promising therapeutic approach for cancer. To date however, the cellular and molecular events governing TLS development have not yet been elucidated. Unravelling these events will be crucial to develop and refine TLS-inducing therapies for cancer treatment.

Glioblastoma (GBM) is a highly aggressive CNS cancer that arises in the supratentorial region of the brain^6^. Standard of care for patients encompasses maximal surgical resection followed by co-administration of temozolomide and fractionated radiotherapy, and adjuvant temozolomide^7^. With this line of treatment, the overall survival of well-selected patients in clinical trials is however still short, with a median of 15-18 months^8^. Non-CNS cancers have benefited from the advent of T cell-reactivating immunotherapies such as checkpoint blockade, which to date have been approved for clinical use in several tumors^9^. However, GBM has proven difficult to treat with checkpoint inhibitors, in both the adjuvant and neo-adjuvant settings^10–14^. The low abundance of T cells within these tumors is likely a contributing factor to their resistance to immunotherapy. We have previously demonstrated that the spontaneous occurrence of TLS in GBM is associated with enhanced T cell infiltration^15^, and that inducing these structures with vascular-targeted AAV-LIGHT therapy promotes survival and anti-tumor responses in murine models of glioma^16^. To date however, the prognostic value of TLS in human GBM remains unknown.

In this study, we analyzed tissues from a cohort of GBM patients, demonstrating that TLS are associated with prolonged survival in this CNS cancer. In addition, we coupled advanced spatial transcriptomics of human GBM tissues with studies in murine models of glioma to resolve the cellular and molecular events leading to TLS formation, elucidating the central role of CD4 T cells in this process.

## Results

### Tertiary lymphoid structures correlate with prolonged survival in glioblastoma patients

To investigate whether TLS presence had prognostic value in patients with high grade astrocytoma, we screened a cohort of 236 patients diagnosed with GBM (IDHwt) or IDH mutant (IDHmut) grade 4 astrocytoma (Cohort 1, Supplementary table 1). FFPE sections obtained from surgical tissues collected before radio-chemotherapy were used to identify lymphoid aggregates through visual assessment of H&E staining. Sequential sections were stained for T cells (CD3), B cells (CD20), endothelial cells (CD34) and nuclei to confirm TLS presence and stratify patients for survival analysis (Figure 1A,B). TLS were defined as tight aggregates of lymphocytes that exhibited a vascular network. The proportion of patients positive for TLS was similar among GBM cases (14.6%) and patients with IDHmut grade 4 astrocytoma (20.8%) (Figure 1C). Survival analysis was performed on patients diagnosed with GBM and included in Cohort 1, who underwent tumor resection followed by standard of care treatment. Inclusion criteria for this analysis comprised (i) confirmed IDHwt status, (ii) access to full survival information and (iii) access to FFPE tissue for TLS assessment, including a total of 188 patients (Supplementary table 1). TLS presence stratified patients into separate prognostic groups, were TLS^+^ patients displayed a significantly prolonged survival (Figure 1D). Altogether, this data indicates that TLS can form in both GBM and IDHmut grade 4 astrocytoma, and that they are associated with improved prognosis in GBM.

**Figure 1.**
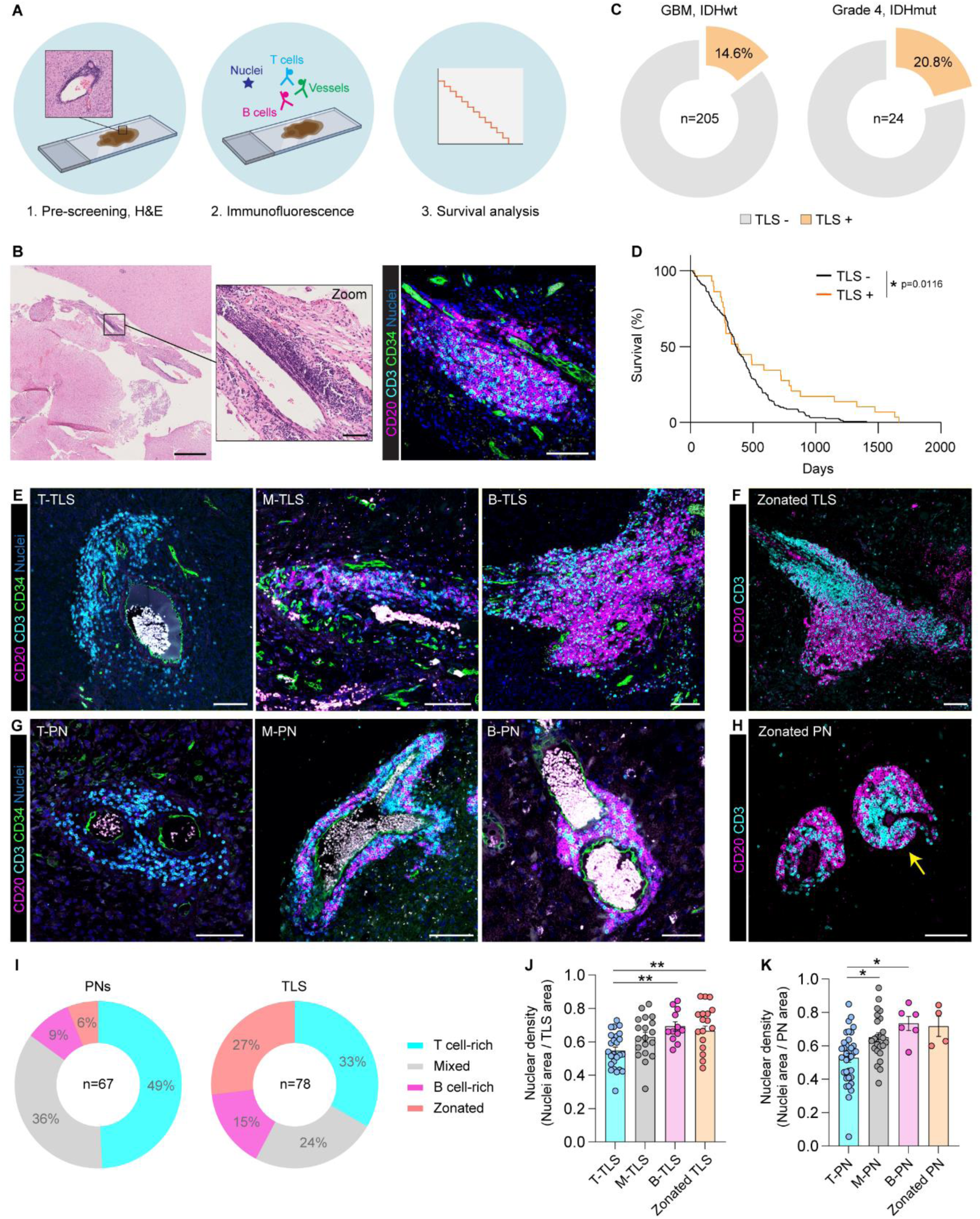
TLS correlate with improved prognosis in glioblastoma, and display variable lymphocytic composition. **(A)** Schematic representation of the experimental layout used to obtain data shown in panels B-K. In brief, sections from large surgical tissues of grade 4 astrocytoma cases (including GBM and IDHmut astrocytomas) were stained with H&E and pre-screened for the presence of lymphoid aggregates. Positive samples were sequentially stained using immunofluorescence (IF) to identify B cells, T cells, endothelial cells and nuclei, and confirm the presence of TLS. Finally, clinical data and staining results were correlated to perform survival analysis. **(B)** Representative images showing a lymphoid aggregate in an H&E-stained section (left panel) and the sequential section stained with IF antibodies for B cells (CD20), T cells (CD3), endothelial cells (CD34) and nuclei (right panel). Zoom panel shows a close-up of the lymphoid aggregate in the H&E-stained section. Scale bar in H&E section: 1 mm. Scale bar in H&E zoom and IF section: 50 µm. **(C)** Pie charts showing the percentage of TLS positive patients in a large GBM cohort (n=205 patients) and a small cohort of grade 4 astrocytoma patients with IDH mutation (n=24 patients). **(D)** Kaplan-Meier survival curve of GBM patients stratified based on TLS presence in surgical tissues. n(TLS-)=159; n(TLS+)=29. Statistics: Log-Rank test. **(E)** Representative IF images of TLS dominated by T cells (T cell-rich TLS, or T-TLS), TLS with a similar representation of B cells and T cells (mixed TLS, or M-TLS) and TLS with a large B cell component (B cell-rich, or B-TLS). **(F)** Representative IF image of a B-TLS with clear zonation of B cells and T cells. **(G)** Representative IF images of PNs dominated by T cells (T cell-rich PNs, or T-PNs), PNs with a similar representation of B cells and T cells (mixed PNs, or M-PNs) and PNs with a large B cell component (B cell-rich, or B-PNs). **(H)** Representative IF image of a B-PN with clear zonation of B cells and T cells. Yellow arrow indicates the zonated PN. Scale bars in (E-H)=100µm. **(I)** Pie charts showing the distribution of T cell-rich, mixed, B cell-rich and zonated structures among TLS and PNs. n(PNs)=67; n(TLS)=78. **(J-K)** Nuclear density in (J) TLS and (K) PNs with different T-to-B cell ratios. n=4-35 aggregates/group. Bar graphs show mean ±SEM. Statistics: one-way ANOVA with Tukey’s correction for multiple comparisons. *p<0.05; **p<0.01.

### Lymphoid aggregates exhibit variable T to B cell proportions and distinct vascular organizations

Immunofluorescence stainings revealed that proportions of T cells and B cells within TLS varied across structures. While some TLS were dominated by T cells and showed very low B cell constituents (T cell-rich TLS, or T-TLS), some showed a similar T-to-B cell content (mixed TLS, or M-TLS) and others exhibited a more pronounced B cell component, while still containing numerous T cells (B cell-rich TLS or B-TLS) (Figure 1E, Supplementary Figure 1A). A proportion of B cell-rich TLS exhibited clear zonation of B cells and T cells (Figure 1F). Interestingly, perivascular aggregates of lymphocytes forming around vessels with characteristically large lumens were also identified (perivascular niches, or PNs), where B cells and T cells remained tightly confined to the perivascular space (Figure 1G,H). Similar to TLS, the T-to-B cell ratio was heterogeneous across PNs (Figure 1G, Supplementary Figure 1B), and some B cell-rich PNs displayed clear B and T cell zonation (Figure 1H). However, while the proportion of B cell-rich/zonated structures was large among TLS (42%), PNs were preferentially T cell-rich and showed little representation of B cell-rich/zonated aggregates (15%) (Figure 1I). Finally, T cell-rich aggregates displayed the lowest nuclear density in both TLS and PN groups (Figure 1J,K), indicating a lesser degree of cell clustering in these structures.

In summary, this data demonstrates that vascular organization and T-to-B cell proportions differ across lymphoid aggregates in GBM tissues, where TLS exhibit a more complex vascular network and a large proportion of B cell-rich structures, while PNs localize around vessels with a large lumen and are preferentially T cell-rich.

### GeoMx spatial analysis identifies key cell types and gene signatures of lymphoid aggregates with different lymphocytic composition

To obtain a better understanding of composition and molecular signatures of TLS and PNs in GBM, lymphoid aggregates present in human GBM tissues were analyzed using the GeoMx spatial transcriptomic platform (Figure 2A). In line with what we previously observed in stained tissues, cell type deconvolution confirmed that T cell-rich structures contained low proportions of B cells, M-TLS had a similar T-to-B cell ratio and B-TLS were largely formed by B cells but still exhibited relevant T cell amounts (Figure 2B). Interestingly, CD4^+^ T cells constituted the largest portion of T lymphocytes in T-PNs, T-TLS and B-TLS, while a 1:1 ratio of CD4^+^ and CD8^+^ T cells was observed in M-TLS (Figure 2B). Among all structures, T-PNs had the most pronounced dendritic cell (DC) signature, while T-TLS exhibited the largest proportion of fibroblasts (Figure 2B). Finally, B-TLS exhibited the lowest fraction of macrophages and, together with M-TLS, displayed gene signatures of plasma cells (PCs) (Figure 2B).

**Figure 2.**
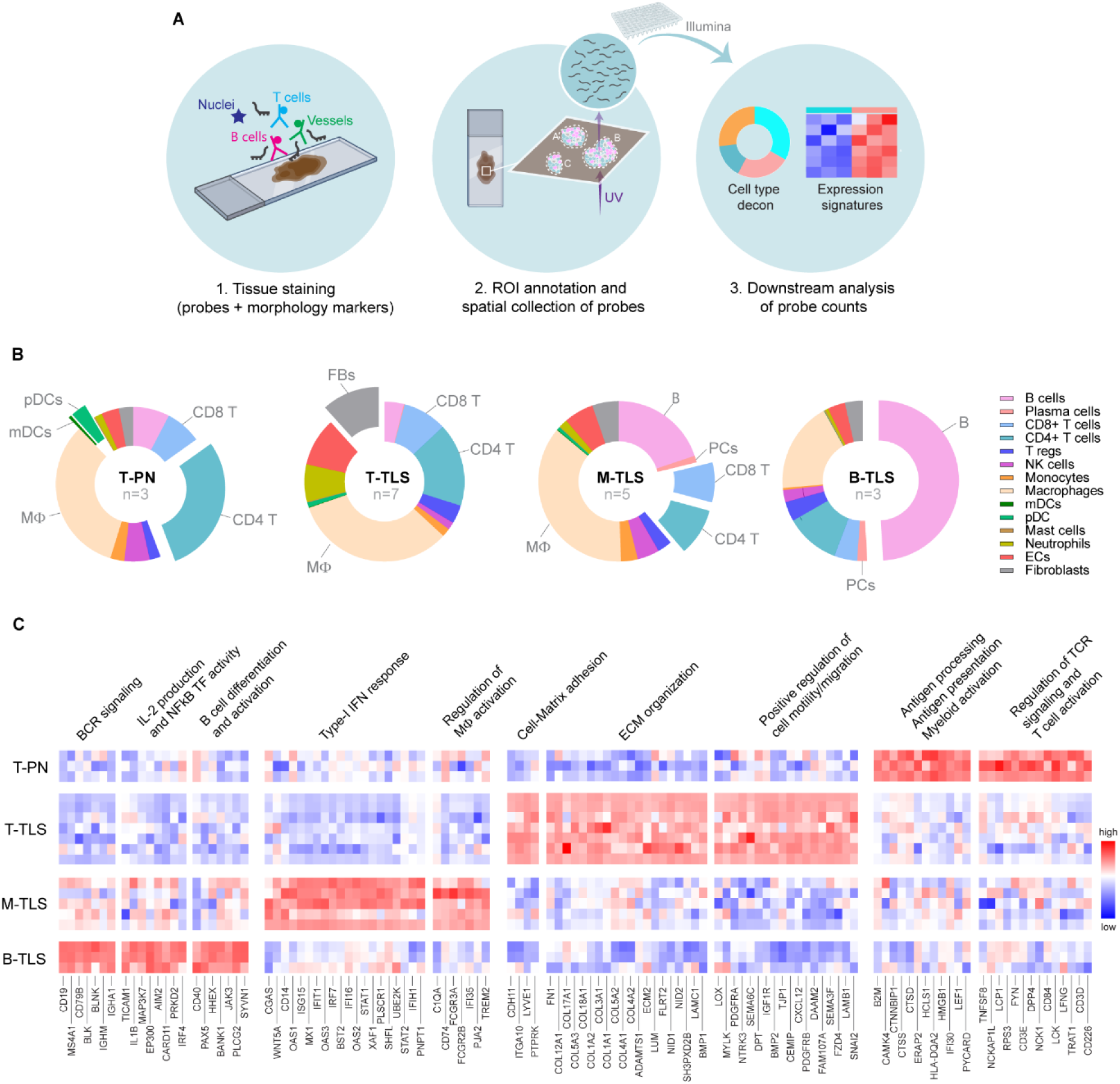
GeoMx spatial analysis identifies cell composition and expression signatures of lymphoid aggregates with distinct T-to-B cell ratios. **(A)** Schematic representation of the GeoMx experimental layout, used to obtain data shown in panels B-C. In brief, FFPE sections of human GBM tissue were co-stained with fluorescent antibodies/dyes targeted to morphological markers to visualize T cells, B cells, endothelial cells and nuclei, and with probes linked to UV-cleavable oligos and targeted to the whole transcriptome. The tissue was scanned and morphological markers were used to identify and annotate lymphoid aggregates and their T-to-B cell composition. Individual aggregates or part of a larger aggregate were selected as regions of interest (ROIs), and UV-cleaved oligos from each ROI were sequenced using the Illumina system. Data were converted into mRNA counts and analyzed to deconvolute cell types and identify gene expression signatures. **(B)** Pie charts displaying the results of cell type deconvolution as the proportions of the indicated cell types within lymphoid aggregates annotated as T-PNs, T-TLS, M-TLS and B-TLS. “n” indicates the number of ROIs. **(C)** Heatmap displaying the relative levels of differentially expressed genes in T-PNs, T-TLS, M-TLS and B-TLS, and the pathways that these genes are enriched within. In (B-C): n(T-PNs)=3; n(T-TLS)=7; n(M-TLS)=5; n(B-TLS)=3.

Next, differential gene expression analysis was coupled with pathway analysis to identify biomarkers of T-PNs, T-TLS, M-TLS and B-TLS (Supplementary table 2), and their related biological processes (Supplementary table 3). T-PNs expressed higher levels of genes involved in myeloid activation, antigen processing/presentation, as well as regulation of T cell receptor (TCR) signaling and T cell activation (Figure 2C). T-TLS were enriched for genes involved in cell-matrix adhesion, extracellular matrix (ECM) organization and positive regulation of cell motility/migration (Figure 2C). Genes involved in type-I interferon (IFN) responses and regulation of macrophage activation were upregulated in M-TLS, while genes required for B cell receptor (BCR) signaling, IL-2 production, NFκB activity as well as B cell differentiation/activation were more highly expressed within B-TLS (Figure 2C).

Altogether, these results show that specific cellular components and key gene signatures characterize lymphoid aggregates with different lymphocytic proportions.

### T cell-rich and B cell-rich TLS represent different stages of TLS assembly

Our GeoMx analysis indicated that T-TLS exhibited minimal B cell content, while B-TLS consistently contained a substantial proportion of T cells. Thus, we hypothesized that TLS with different T-to-B cell ratios could represent stages of TLS assembly, where T cells are recruited first and B cells later. To test this hypothesis, we performed a longitudinal study in a murine glioma model, where we orthotopically injected mice with CT-2A glioma cells and collected tumor-bearing brains at three different time points, representing early and late stages of tumor development. Sections from brains collected at each time point were screened for TLS and PNs using immunofluorescence (Figure 3A). In line with data from human GBM tissues, TLS exhibited lymphocytic clustering around a more complex vascular network, while PNs were characterized by B cells and T cells tightly wrapped around a main vessel with a large lumen (Figure 3B). TLS at earlier time points (day 10 and day 17 post-tumor implantation) were largely T cell-rich, while at the survival endpoint a large proportion of B cell-rich TLS was observed (Figure 3C,D). In line with this, the CD3-to-B220 ratio within the TLS dropped significantly between day 17 and the survival endpoint (Figure 3E). Similar to what was observed in human GBM, PNs were largely T cell-rich at all timepoints (Figure 3C,D), and no significant differences were detected in their CD3-to-B220 ratio over time (Figure 3F). No significant differences were identified in TLS or PN numbers over time (Supplementary Figure 2A,B). Interestingly, in human tissue instances were observed where T-PNs forming around large vessels were surrounded by additional areas of dense lymphocyte clustering, associated with smaller vasculature (Supplementary Figure 2C). To understand whether these vascular beds were connected or independent, this instance was investigated in the CT-2A murine glioma model using thicker tissue sections. Here we found that PNs exhibited smaller branches, along which further lymphocytic recruitment was observed (Figure 3G). This suggests that the recruitment of lymphocytes starts within T-PNs around a main large vessel, and progresses into the reorganization of these cells around a more complex vascular network to potentially form TLS.

**Figure 3.**
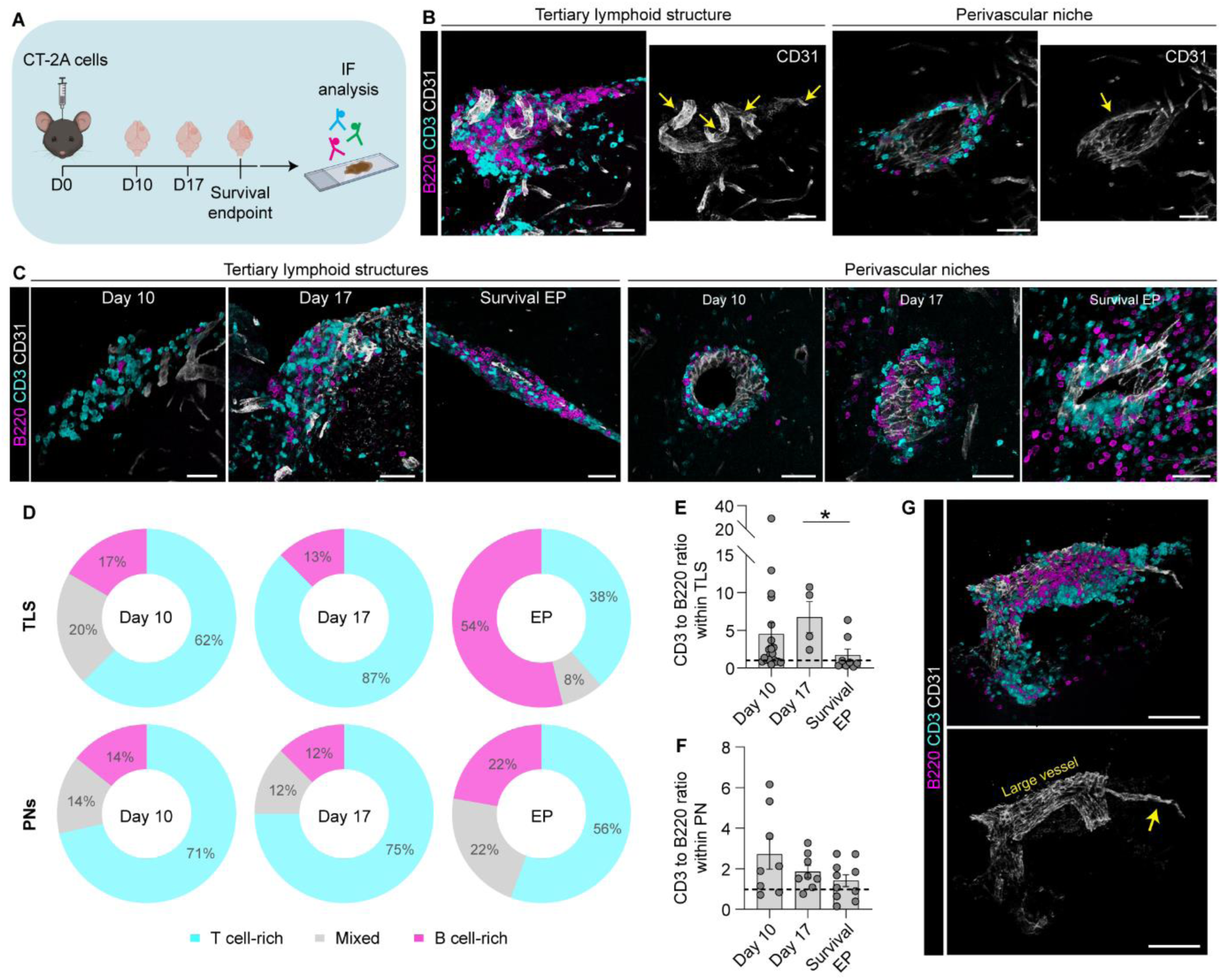
T cell recruitment precedes that of B cells during TLS assembly. **(A)** Schematic representation of the experimental layout used to obtain data shown in panels B-G. In brief, immunocompetent mice were orthotopically injected with syngeneic CT-2A glioma cells, and glioma-bearing brains were collected at day 10 and day 17 post-tumor implantation, as well as at the survival endpoint. Brain sections were stained for B cells (B220), T cells (CD3) and endothelial cells (CD31) using immunofluorescence (IF) to visualize TLS. n=5-10 mice/time point. **(B)** Representative IF images of TLS and PNs present in CT-2A glioma-bearing brains. Scale bars: 50µm. **(C)** Representative IF images of TLS and PNs at the indicated time points post-tumor implantation. Scale bars: 50µm. **(D)** Percentages of T cell-rich, mixed and B cell-rich TLS and PNs at the indicated time points post-tumor implantation. **(E,F)** CD3-to-B220 ratio (CD3+ area / B220+ area) within (E) TLS and (F) PNs at the indicated time points post-tumor implantation. Dots represent individual aggregates. n(TLS)=4-20/group; n(PNs)=8-10/group **(G)** Representative IF image of a large vessel surrounded by a PN, displaying smaller branching along which further lymphocytic recruitment is observed. Scale bars: 100µm.

In summary, this data suggests that T-PNs could represent the seeding events preceding TLS formation in GBM. Moreover, these results demonstrate that T cells and B cells are recruited in a temporally distinct manner during TLS assembly, indicating that T-TLS and B-TLS represent earlier and later stages of TLS assembly, respectively.

### Single-cell spatial analysis identifies lymph node-like spatial organization of immune and stromal cells within TLS

To further dissect TLS characteristics in GBM, sections from cryopreserved surgical GBM tissues were analyzed using Padlock probe-based targeted *in situ* sequencing (Supplementary Figure 3A, Supplementary table 4), and the distribution of immune-related mRNAs within TLS^+^ tissue areas was investigated. Lymphoid aggregates were identified by visualizing the location of T cell and B cell-related mRNAs such as CD3D, CD3E and MS4A1 (Figure 4A). The distribution of B cell-related mRNAs like CD79A, CD79B and CD40 followed the one of MS4A1, while TRBC1 and TRBC2 transcripts followed the distribution of CD3D and CD3E (Figure 4A-D). Among the transcripts that were enriched in the TLS, we identified lymph node-related genes such as CXCR4, CCL5, CD7 and GPR183. CXCR4 was evenly distributed across the whole structure (Figure 4E), suggesting expression by both B cells and T cells, while CCL5 and CD7 preferentially localized within the T cell zone (Figure 4F,G). Finally, GPR183 expression was more concentrated in the B cell zone (Figure 4H).

**Figure 4.**
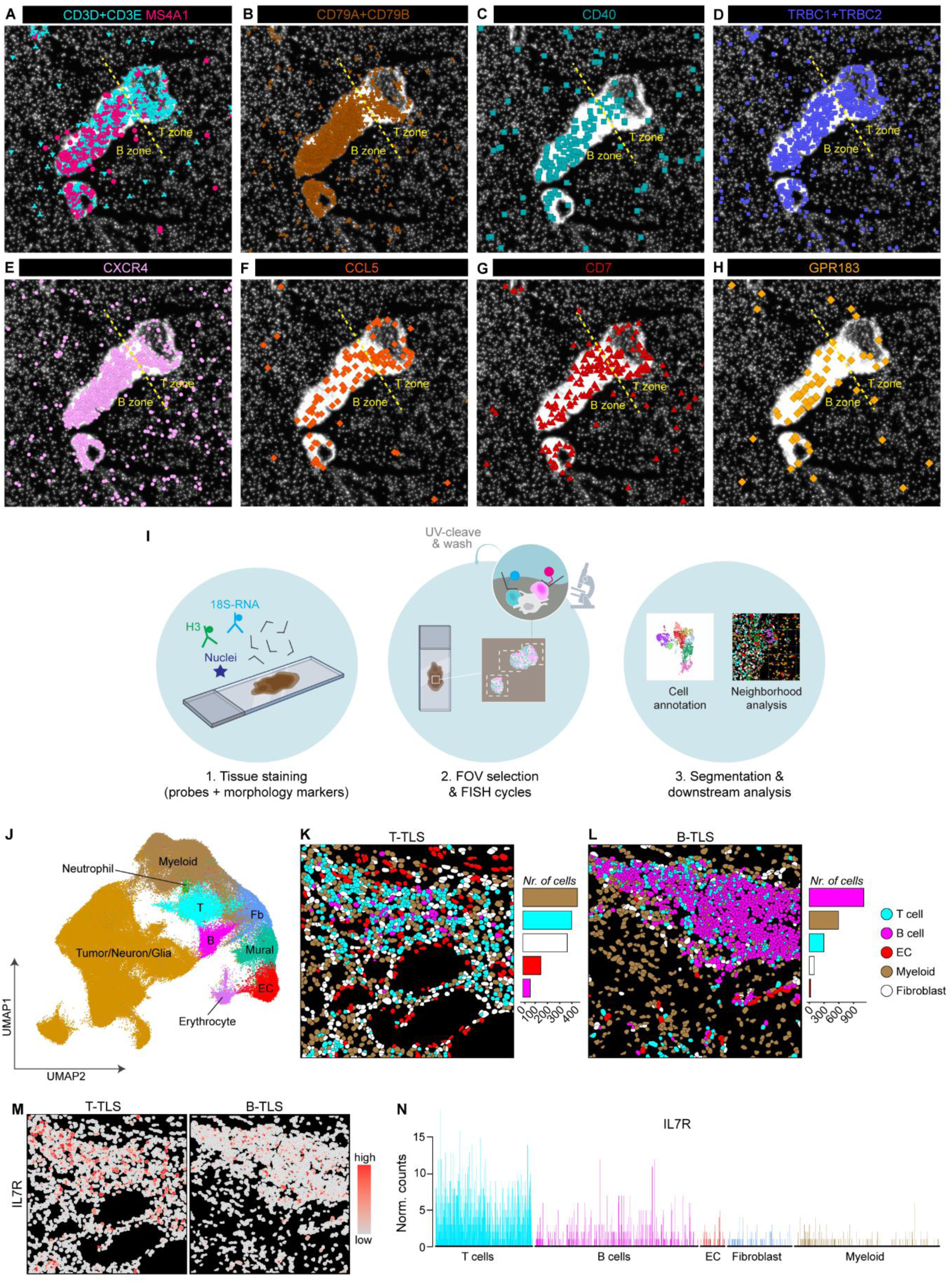
Targeted *in situ* sequencing and CosMx single-cell spatial analysis reveal cell-specific organization and gene expression within TLS. In this figure, data in panels A-H were produced using targeted Padlock probe-based *in situ* sequencing, and data in panels I-M were produced using CosMx single-cell spatial analysis. **(A)** Representative image of a zonated TLS identified in human GBM tissue using targeted Padlock probe-based *in situ* sequencing. The image shows highly clustered signals from CD3D+CD3E (T cell) and MS4A1 (B cell) transcripts, segregated into clear zones. **(B-H)** Location of the following transcripts within the zonated TLS shown in panel A: (B) CD79A and CD79B; (C) CD40; (D) TCRB1+TCRB2; (E) CXCR4; (F) CCL5; (G) CD7 and (H) GPR183. In (A-H): spatial coordinates of transcripts are indicated by colored symbols. White = nuclei. Dotted yellow line separates B versus T cell zone of the shown TLS. **(I)** Schematic representation of the layout used for the CosMx single-cell spatial transcriptomics assay. In brief, FFPE sections of human GBM tissue were co-stained with fluorescent antibodies/dyes to visualize the nuclear (H3) versus cytoplasmic (18S RNA) cell fractions, as well as with *in situ* hybridization (ISH) probes targeted against 1030 mRNAs. Fields of view (FOVs) were selected, and probes were detected using complementary reporter probes tagged with fluorophores. Probe signal was detected within each FOV using cycles of fluorescence imaging, and probe location on the tissue was recorded using tissue coordinates. Cells were annotated based on clustering analysis and gene expression profiles. Annotated cells were plotted back onto their tissue coordinates to visualize their location. **(J)** UMAP of main cell clusters annotated in GBM tissue sections using CosMx single-cell spatial analysis (n=4 patients). T= T cell; B= B cell, Fb= fibroblast. **(K,L)** Spatial organization of T cells, B cells, endothelial cells (ECs), myeloid cells and fibroblasts in (J) T cell-rich TLS (T-TLS) and (D) B cell-rich TLS (B-TLS). Images show representative FOVs containing T-TLS and B-TLS. Bar graphs to the right of each image display the number of cells within the shown FOVs, for each plotted cell type. **(M,N)** Expression of IL7R in single cells within T-TLS and B-TLS visualized as (L) an expression heatmap or (B) a bar graph displaying normalized counts of IL7R in each cell within the shown FOVs.

To investigate the organization and gene expression of immune and stromal cells within TLS at a single-cell resolution, FFPE sections collected from human GBM tissues were analyzed using the CosMx single-cell spatial transcriptomic platform (Figure 4I, Supplementary table 5). Single cells were clustered based on similarities in gene expression (Supplementary Figure 4A) and main cell types were annotated using expression scores (Figure 4J, Supplementary Figure 4B-J, Supplementary table 6). Visualization of the annotated cell types at their original tissue coordinates allowed for the identification of T-TLS and B-TLS (Figure 4K,L). T-TLS contained large numbers of myeloid cells and fibroblasts, which were present within the structure in close proximity of the T cells (Figure 4K), reminiscent of the organization of fibroblastic reticular cells (FRCs) within the T cell zone of a lymph node. In B-TLS instead, T cells were scattered across a tight B cell core, while myeloid cells and fibroblasts were mostly located in the periphery (Figure 4L), organizing similarly to a B cell follicle. Next, we studied the expression of lymph node-related genes in the main cell populations found within TLS. In line with our *in situ* sequencing results, CCL5 was predominantly expressed by T cells (Supplementary Figure 4K,L), while the expression of CXCR4 was enriched in both T cells and B cells (Supplementary Figure 4M,N). Interestingly, the expression of CXCR4’s ligand CXCL12 was enriched in myeloid cells and fibroblasts (Supplementary Figure 4O,P), indicating a possible interplay between these cells and lymphocytes within TLS. Interestingly, IL7R, a marker of lymphoid tissue-inducer (LTi) cells, was preferentially expressed by T cells within TLS (Figure 4M,N), suggesting a potential LTi role for this cell population in GBM-associated TLS.

In summary, this data suggests that the organization and gene expression of immune and stromal cells in T-TLS and B-TLS resembles the one of T cell zones and B cell follicles of a lymph node.

### Availability of MMP2^+^ meningeal fibroblasts correlates with the organization of lymphocytes beyond the perivascular space

Next, we sought to further investigate the connection between PNs and TLS. Notably, lymphoid aggregates organized as TLS were more likely to be present in proximity of meningeal-rich areas of the brain, while PN-like aggregates were more often found in distal brain areas (Figure 5A). To investigate which discriminating factor between these two brain locations correlated with the ability of perivascular lymphoid aggregates to re-organize as TLS, we analyzed the cellular composition of meningeal versus intratumoral PNs (Figure 5B,C). Cellular components in meningeal T-PNs were similar to those of T-TLS and included a large proportion of fibroblasts (Figure 5B), while intratumoral T-PNs lacked fibroblastic presence (Figure 5C). A striking difference in collagen IV organization was also observed when comparing intratumoral PNs and meningeal TLS with similar lymphocytic composition. Indeed, intratumoral PNs were encapsulated by a surrounding collagen IV^+^ basement membrane, preventing lymphocytes from exiting the perivascular space (Figure 5D). Conversely, lymphocytes within meningeal TLS were located beyond the perivascular collagen IV barrier. In line with this, fibroblasts present in meningeal PNs or TLS expressed high levels of MMP2, the metalloproteinase responsible for cleaving collagen IV fibers (Figure 5E). Furthermore, spatially-resolved interaction analysis identified that MMP2^+^ fibroblasts were consistently located in close proximity of endothelial cells (ECs) within meningeal aggregates (Figure 5F,G), while no such interactions were identified within intratumoral PNs.

**Figure 5.**
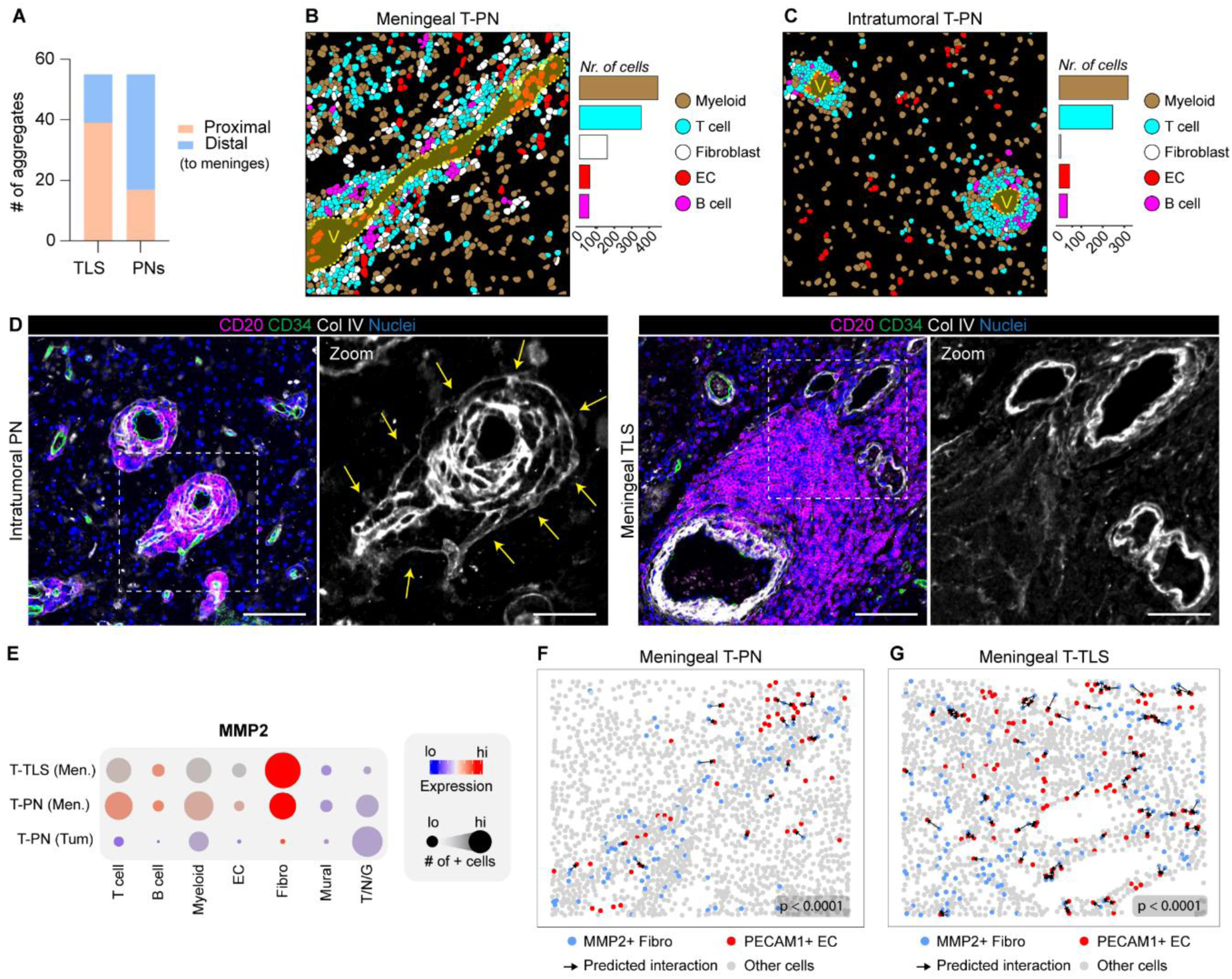
Availability of meningeal MMP2^+^ fibroblasts correlates with disruption of perivascular collagen IV capsule and TLS-like arrangement of lymphocytes. Panels A and D show data obtained via immunofluorescence analysis of human grade 4 astrocytoma tissues (including cases of GBM and IDHmut grade 4 astrocytoma). The remaining panels show spatial analysis of CosMx data from human GBM tissues. **(A)** Quantification of the number of TLS and PNs found proximally (proximal) or distally (distal) to meningeal-rich areas of the brain. n=56 aggregates/group. **(B,C)** Spatial organization of T cells, B cells, endothelial cells (ECs), myeloid cells and fibroblasts in (B) meningeal and (C) intratumoral T-PNs. Images show representative fields of view (FOVs) containing meningeal and intratumoral T-PNs. Lumens of vessels within each PN are highlighted in yellow, and marked as “V”. Bar graphs to the right of each image display the number of cells within the shown FOVs, for each plotted cell type. **(D)** Representative immunofluorescence images of the distribution of collagen IV in intratumoral PNs versus meningeal TLS. Dotted squares indicate areas shown in the “Zoom” panels to the right of each image. Yellow arrows indicate collagen IV capsule surrounding perivascular lymphocytes. Main scale bars: 100µm. Zoom scale bars: 50µm. **(E)** Bubble plot showing the expression of MMP2 in the indicated cell types across representative FOVs containing meningeal T-TLS, meningeal T-PNs and intratumoral T-PNs (shown in Figure 4K, 5B and 5C). Bubble’s size indicates the number of MMP2^+^ cells within the FOV, for each cell type. Color scale indicates average MMP2 expression levels for each cell type. EC= endothelial cell, T/N/G= Tumor/Neuron/Glia. **(F,G)** Interaction plots showing significant predicted interactions between MMP2^+^ fibroblast and PECAM1^+^ ECs in (F) meningeal T-PNs and (G) meningeal T-TLS. Black arrows indicate predicted interactions; lo=low; hi=high; # of + cells = number of MMP2^+^ cells. Distribution of major cell types in the shown FOVs can be found in Figure 4K and Figure 5B.

Altogether, this data suggests that perivascular recruitment of immune cell types present within lymphoid aggregates can occur similarly within the tumor and in proximity of the meninges. However, the availability of MMP2^+^ meningeal fibroblasts correlates with the ability of these aggregates to organize as TLS (with lymphocytes arranging outside of the perivascular space) rather than remaining confined by perivascular collagen IV fibers.

### CD4^+^ T cells exhibit molecular characteristics and interactions typical of LTi cells

Having established that TLS assembly in GBM begins with the perivascular recruitment of T cells and with their reorganization into T-TLS, we sought to investigate which molecular events characterized these early stages of TLS development. To answer this question, we used our CosMx dataset to perform an intra-patient analysis aimed at identifying molecules and interactions characteristic of T-PNs and T-TLS. From our GeoMx and CosMx analyses, we learned that CD4 T cells were the major lymphocytic component of T-PNs (Figure 2B), and that T cells within TLS expressed high levels of the LTi marker IL7R (Figure 4M,N). In addition, CD4 T cells within PNs exhibited the highest levels of CCR7 (Figure 6A, Supplementary table 7), a crucial receptor for the initial accumulation of LTi cells during lymphoid development^17^. In line with this, predicted interaction between CCR7^+^ CD4 T cells and multiple cell types expressing CCR7’s ligands were identified within T-PNs, including those with CCL19^+^ ECs (Figure 6B) and CCL19^+^/CCL21^+^ myeloid cells (Figure 6C and Supplementary Figure 5A). In addition, CD4 T cells exhibited higher expression of IL7R than CD8 T cells (Figure 6D, Supplementary table 8), making them more likely LTi candidates. IL7R^+^ CD4 T cells were also predicted to interact with IL7^+^ myeloid cells and IL7^+^ fibroblasts within T-TLS (Figure 6E,F), an interaction necessary for the stimulation and sustenance of LTi cells throughout lymphoid development^18,19^. CD4 T cells involved in these interactions also expressed genes found downstream of the IL7-IL7R signaling cascade (Figure 6G,H). Similar results were found for the aforementioned interactions involving CCR7^+^ CD4 T cells (Supplementary figure 5B,C). Finally, myeloid cells within T-TLS, but not ECs or fibroblasts, showed high expression of VCAM1 (Figure 6I), a marker of activated lymphoid tissue-organizer (LTo) cells. In line with this, high numbers of predicted interactions of CXCL12^+^ myeloid cells with both CXCR4^+^ CD4 T cells and CXCR4^+^ B cells were identified within T-TLS (Figure 6J,K), suggesting that myeloid cells harbored the most prominent LTo potential.

**Figure 6.**
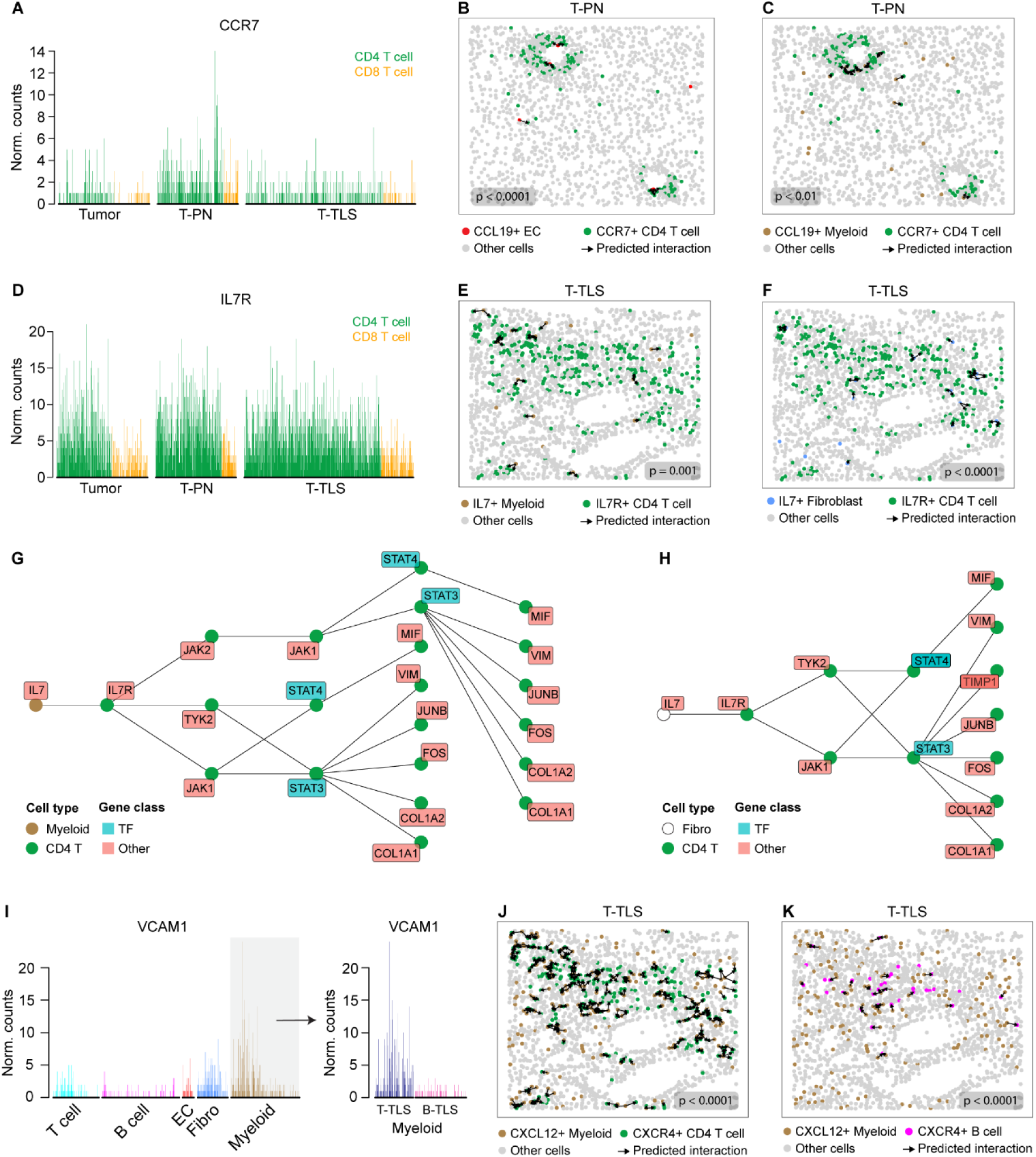
Molecular interactions between CD4 T cells and immune/stromal cells within T cell-rich aggregates are reminiscent of LTi-LTo cross-talk during lymphoid tissue development. Panels A-H show results from intra-patient spatial analysis performed in GBM tissue from patient #240, using CosMx data. Distribution of major cell types in the FOV shown in panels B,C can be found in Figure 5C. Distribution of major cell types in the FOV shown panels E,F,J and K can be found in Figure 4K. **(A)** Single-cell expression of CCR7 in CD4^+^ T cells and CD8^+^ T cells located within T-PNs, T-TLS and tumor area. **(B,C)** Interaction plots showing significant predicted interactions between CCR7^+^ CD4 T cells and (B) CCL19^+^ endothelial cells (ECs) or (C) CCL19^+^ myeloid cells in T-PNs. **(D)** Single-cell expression of IL7R in CD4 T cells and CD8^+^ T cells located within T-PNs, T-TLS and tumor area. **(E,F)** Interaction plots showing significant predicted interactions between IL7R^+^ CD4 T cells and (E) IL7^+^ myeloid cells or (C) IL7^+^ fibroblasts in T-TLS. **(G,H)** Pathway-related results of the interaction analysis displayed in panels (E,F), showing that IL7R^+^ CD4 T cells that were predicted to interact with either (G) IL7^+^ myeloid cells or (H) IL7^+^ fibroblasts express genes located downstream of the IL7-IL7R signaling cascade. TF= transcription factor. **(I)** Single-cell expression of VCAM1 across the indicated cell types, within the TLS FOVs shown in Figure 4K,L. Grey box indicates bimodal distribution of VCAM1 expression in the myeloid cell population, related to differences in T-TLS vs B-TLS. **(J,K)** Interaction plots showing significant predicted interactions between CXCL12^+^ myeloid cells and (J) CXCR4^+^ CD4 T cells or (K) CXCR4^+^ B cells in T-TLS. In (B,C,E,F,J,K) black arrows indicate predicted interactions.

In summary, this data indicates that CD4 T cells found within PNs and TLS exhibit molecular characteristics typical of LTi cells, and are involved in a variety of interactions with immune and stromal cells displaying LTo-like features.

### IL7R^+^ CCR7^+^ CXCR4^+^ LTB^+^ LTi-like Th1 cells are enriched within TLS

To investigate whether CD4 T cells are necessary for the formation of TLS in GBM, we depleted this population in CT-2A glioma-bearing mice and investigated TLS presence via immunofluorescence analysis of brain sections at the survival endpoint (Figure 7A). Strikingly, depletion of CD4 T cells abrogated TLS formation (Figure 7B,C), indicating that a subpopulation of helper T cells is necessary to initiate this process. To identify which CD4 subpopulation displayed LTi potential in human GBM, we used in-house single-cell sequencing data from CD4 T cells infiltrating human high-grade gliomas (Supplementary Figure 6) to identify relevant genes that could be interrogated in our CosMx dataset (see Methods section for details). Next, we performed a trajectory analysis of CD4 T cells present in tissues included in our CosMx cohort, and annotated each T helper (Th) state based on gene expression (Figure 7D, Supplementary table 9). Six main CD4 T cell states were identified (Figure 7E). Th17 inflammatory (Th17_inf_) cells expressed high levels of IL17RE, IL1R1 and COL5A3, with low expression of checkpoint molecules such as PDCD1, CTLA4 and TIGIT, and intermediate levels of IFNG (Supplementary Figure 7). Th17 inhibitory (Th17_inh_) cells expressed high levels of IL17A, IL17RB, IL1R1 and COL5A3, intermediate levels of PDCD1, CTLA4 and TIGIT and low levels of IFNG (Supplementary Figure 7). Cells in a Th1-Th17 transitory state (Th1-Th17) expressed intermediate levels of both Th1 (e.g. STAT4, GZMK) and Th17 (e.g. IL17A, IL17RB) signature molecules (Supplementary Figure 7). T regulatory cells (Tregs) expressed the highest levels of FOXP3, IL2RA, and inhibitory molecules such as PDCD1, CTLA4 and TIGIT (Supplementary Figure 7). Finally, two states with Th1 characteristics were identified, namely Th1 resting (Th1_rest_) and Th1 activated (Th1_act_) cells, both expressing the highest levels of signature Th1 markers (such as TBX21, STAT4, GZMK, CCL5 and CXCR3) and of the LTi marker IL7R (Figure 7F, Supplementary Figure 7). Compared to Th1_act_ cells, Th1_rest_ cells exhibited lower levels of functional molecules such as IFNG, GZMA and GZMH, higher expression of the early activation marker CD69, but low levels of effector/late activation markers such as CD44, CD28 and checkpoint molecules. They also displayed higher levels of the chemotactic receptors CCR7 and CXCR4 (Figure 7F), suggesting that they had been recently recruited into the tissue. Th1_act_ cells instead displayed intermediate levels of CCR7 and CXCR4, but higher expression of both functional and effector/late activation markers (Figure 7F). Cells in both Th1 states expressed the LTi molecule LTB, with Th1_act_ cells displaying the highest levels (Figure 7F), in line with their enhanced activation status. Both Th1 subpopulations also expressed high levels of CD40LG. Importantly, Th1 cells were specifically enriched within PNs and TLS compared to the tumor area, which was instead preferentially inhabited by suppressive CD4 populations, such as Tregs and Th17_inh_ cells (Figure 7G,H, Supplementary Figure 8A,B). Moreover, the proportion of Th1 cells with LTi potential increased across the developmental timeline of TLS assembly, with the highest proportion found within B-TLS (Figure 7G and Supplementary Figure 8C).

**Figure 7.**
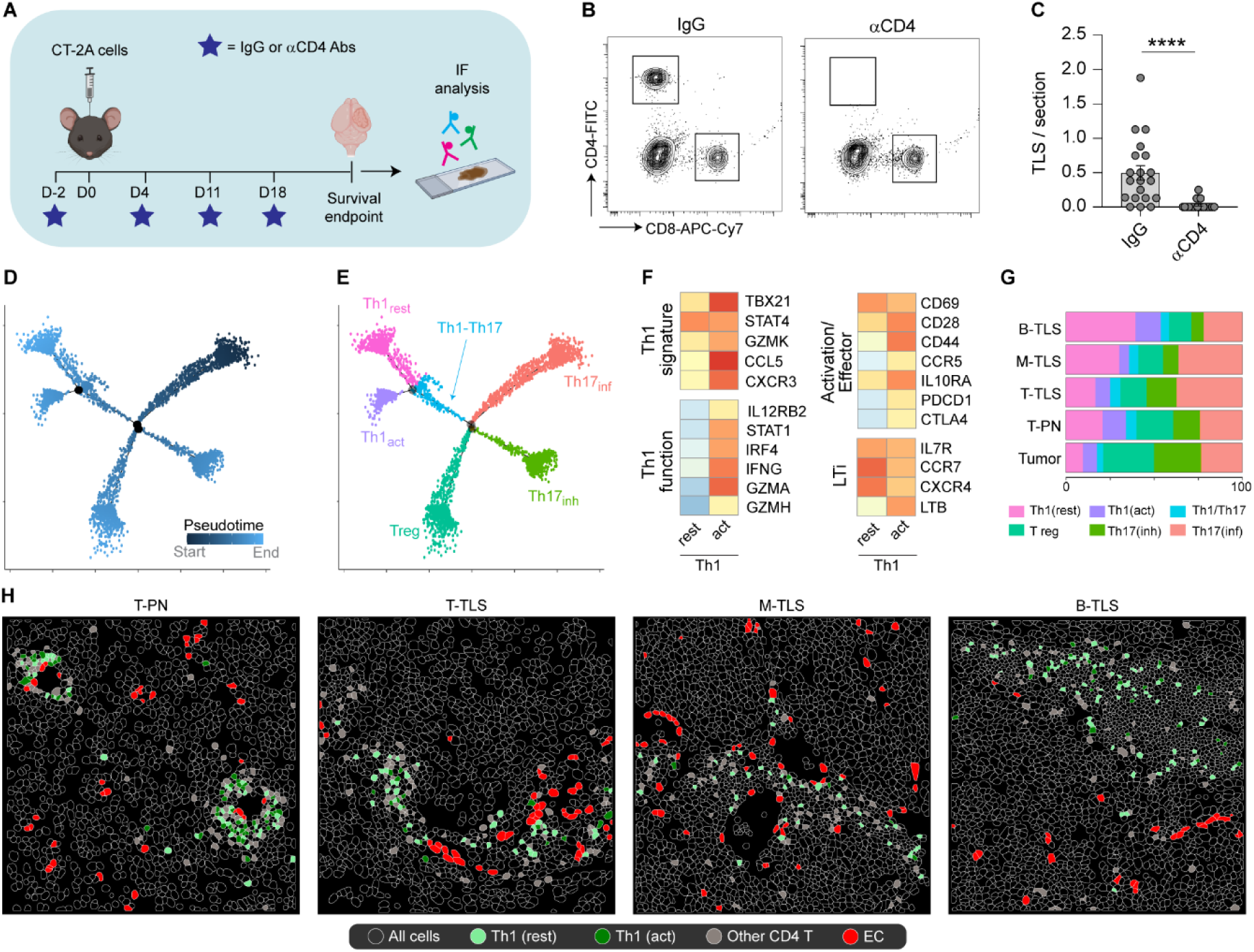
IL7R^+^CCR7^+^LTB^+^ Th1 cells with lymphoid tissue-inducing (LTi) potential preferentially localize within PNs and TLS in GBM. Panels (A-C) show mouse-related data. Panels (D-I) show data obtained via CosMx spatial analysis of human GBM tissues. Panels (J and K) show data obtained via targeted Padlock probe-based *in situ* sequencing of human GBM tissue. **(A)** Schematic representation of the experimental layout used to obtain data shown in panels B-C. In brief, immunocompetent mice received either IgG or αCD4 antibodies on day -2 before being orthotopically injected with syngeneic CT-2A glioma cells on day 0. IgG or αCD4 antibodies were administered also on days 4, 11 and 18 post-tumor implantation. Brains were collected at the survival endpoint and processed for immunofluorescence analysis to quantify TLS. n=20 mice/group. **(B)** Representative FACS plots showing the absence of a CD4^+^ T cell population in the blood of αCD4-treated mice. **(C)** Number of TLS per section in the indicated groups. n=20 mice/group. Statistics: Mann-Whitney test; ****p<0.0001. **(E)** Distribution of CD4 T cells from GBM tissues (CosMx cohort) along a pseudotime-based trajectory. **(F)** Annotation of six main CD4 states/subpopulations along the trajectory shown in panel (E). **(G)** Heatmap displaying the relative expression levels of markers related to Th1 signature, Th1 function, Activation/Effector molecules and LTi molecules in the Th1(rest) and Th1(act) subpopulations. **(G)** Proportions of each CD4 subpopulation in tumor, T-PNs, T-TLS, M-TLS and B-TLS. **(H)** Representative images of the spatial distribution of the Th1(rest) and (Th1act) subpopulations in T-PNs, T-TLS, M-TLS and B-TLS. Distribution of major cell types in the shown FOVs can be found in Figure 5C, Supplementary Figure 8A and Figure 4L.

Altogether, this data indicates that a subpopulation of pro-inflammatory IL7R^+^CCR7^+^CXCR4^+^ Th1 cells is recruited and maintained specifically at sites of TLS formation in GBM tissue. Once activated, these cells can undertake the role of LTi cells, by producing high levels of the LTi molecule LTB.

## Discussion

To date, it is accepted that the presence of TLS in proximity of tumor tissues is associated with positive outcomes in several non-CNS cancer types, both in terms of survival and responses to immunotherapy^20–23^. While we have previously demonstrated that TLS correlate with increased T cell abundance in GBM tissues^15^, before this study the prognostic relevance of these structures in CNS cancers was unknown. Here, a large cohort of GBM and IDHmut grade 4 astrocytoma patients was screened for TLS, where TLS were defined as tight clusters of lymphocytes forming around a network of blood vessels. Approximately 15% of IDHwt GBM patients were found to be TLS^+^. While all samples in this cohort were large sections of surgically-resected tissues, not all samples included parts of the meningeal layer, which is the preferential location of TLS formation in murine and human GBM^15,16^. Thus, some TLS^+^ patients could have been missed, which would be in line with the higher proportion of TLS^+^ patients that we previously observed in a smaller GBM cohort, where meningeal tissue was available for all investigated samples^15^. Nevertheless, stratification of patients based on TLS assessment in the current GBM cohort revealed that these structures correlate with prolonged survival in patients that received standard-of-care treatment. This is a promising result, which warrants more investigation into the use of TLS as clinical biomarker of improved prognosis in GBM. Given their association with increased intratumoral T cell infiltration^15^, TLS presence could also be used to select GBM patients that may better respond to T cell-reactivating immunotherapies.

Studies in pre-clinical cancer models have shown that delivering inducer molecules such as lymphotoxins or LIGHT to the tumor microenvironment can induce TLS, prolong survival and boost anti-tumor immune responses^24^. Importantly, we have shown that this can be achieved in murine models of glioma^16^, indicating that TLS-inducing therapies hold promise for the treatment of aggressive CNS tumors. However, the steps and mechanisms through which TLS formation is initiated remain largely unclear, limiting our ability to enhance TLS-inducing therapeutics. To date, most studies of TLS in human cancer have relied on immunohistochemical stainings of sequential tissue sections, or multiplex immunostainings including less than ten parameters^25–27^. These studies have been important in elucidating that TLS exhibit different maturation stages which have an impact on patients’ prognosis, with TLS that resemble secondary follicles being associated with the most positive outcomes^4,5^. Nevertheless, higher resolution studies are needed to unravel the molecular and cellular events involved in initiating the development of these structures, which will be central to the advancement and refinement of treatments aimed at modulating TLS. Here, we dissected the steps leading to TLS formation in GBM by combining advanced spatial transcriptomic techniques with studies in murine models, demonstrating that the assembly of T cell-rich aggregates is a prerequisite for the development of canonical B cell-rich TLS (B-TLS).

We found that TLS development in GBM begins with the recruitment of CD4 T cells around large, activated vessels expressing T cell-recruiting chemokines such as CCL19, leading to the formation of T cell-rich perivascular niches (PNs). PNs were tightly encapsulated by collagen IV fibers, preventing lymphocytes from exiting the perivascular space. This is in line with the profoundly abnormal morphology of the GBM vasculature, which is characterized by enhanced deposition of extracellular matrix (ECM) proteins such as collagen IV^28^. Interestingly, the availability of MMP2^+^ meningeal fibroblasts correlated with the disruption of this capsule and the organization of lymphocytes around a more complex vascular network, to form T cell-rich aggregates reminiscent of TLS (T-TLS). In line with this, T-TLS exhibited enriched expression of genes associated with ECM reorganization and positive regulation of cell migration, indicative of the rearrangement of cells and matrixes within these structures. Aggregates forming distally from the meninges remained more commonly confined within perivascular collagen IV fibers. This is in accordance with the physiological organization of fibroblasts in the brain, which is largely limited to the meninges and Virchow-Robin spaces. These findings shed light on why TLS are preferentially found in close proximity of meningeal tissue in GBM^15^.

T cell-rich structures exhibited a lower nuclear density than B cell-rich structures in human GBM tissues, suggesting that perivascular T cell clustering events could correspond to the initial step of TLS nucleation, when lymphocytes begin to aggregate. In line with this hypothesis, we demonstrated that the assembly of T cell-rich aggregates temporally preceded that of TLS with a defined B cell core in murine glioma. Moreover, in human GBM tissues CD4 T cells present within T cell-rich PNs exhibited receptor-ligand interactions characteristic of early lymphoid recruitment events^17^, including those between CCR7^+^ CD4 T cells and CCL21^+^/CCL19^+^ myeloid cells. The IL-7/IL-7R axis is also critical during lymph node formation, in which the IL-7 produced by LTo cells signals IL-7R^+^ LTi cells to kick-start lymphoid development^29^. Interactions between IL7R^+^ CD4 T cells and IL7^+^ myeloid cells or fibroblasts were observed within T-TLS, reminiscent of events leading to LTi stimulation and activation during lymphoid development^18,19^. Moreover, depletion of CD4 T cells in murine glioma abolished TLS formation, underscoring the importance of CD4 T cell-related events for TLS assembly.

In human GBM, two subpopulations of pro-inflammatory Th1 CD4 T cells with LTi potential were preferentially found within the TLS. These included (1) freshly recruited IL7R^+^CCR7^hi^CXCR4^hi^ CD44^low^ cells, exhibiting a resting profile with low expression of functional molecules, and (2) their activated IL7R^+^CCR7^int^CXCR4^int^CD44^hi^ counterpart, expressing increased levels of interferon-γ and of the TLS-inducing cytokine lymphotoxin-β. Both Th1 subpopulations expressed the co-stimulatory molecules CD28 and CD40LG, thus being equipped to license CD40^+^ DCs for cross-presentation, an important step for the priming and activation of tumor-targeted CD8 T cells^30^. Notably, pro-inflammatory Th1 cells were rarely found outside of lymphoid aggregates or within the tumor area, which was largely inhabited by immunosuppressive CD4 T cells such as Tregs or inhibitory Th17 cells. This indicates that TLS represent immune niches that have the ability to sustain pro-inflammatory phenotypes of T helper cells in GBM.

In summary, our work identifies a correlation between TLS presence and improved survival in human GBM, highlighting their potential as clinical parameters for patient stratification. Moreover, we hereby define the cellular and molecular events leading to TLS assembly in GBM. We propose that pro-inflammatory IL7R^+^ Th1 lymphocytes with LTi potential are recruited to TLS nucleation sites through activated vessels. Here, they can be stimulated by IL7^+^ immune and stromal cells, to initiate the process of TLS assembly via the production of inducer molecules such as LTB. Immune and stromal cells contribute to further T and B cell recruitment within these structures via the CCR7-CCL19/CCL21 and CXCR4-CXCL12 chemotactic axes. Importantly, we also observe that in environments abundant in MMP2^+^ fibroblasts, such as the meninges, lymphocytes are able to organize into TLS outside of the collage IV-coated perivascular space. Conversely, in regions of the brain lacking fibroblasts such as the parenchyma, the collagen IV barrier remains intact, confining lymphocytes to the perivascular space. Altogether, these findings shed light on the cellular and molecular events leading to TLS assembly in GBM, highlighting a central role for CD4 T cells and paving the way for the development of improved TLS-modulating therapies for cancer.

## Methods

### Patient material

A retrospective cohort of 236 cases of grade 4 glioma was assembled (Supplementary table 1, Cohort 1). This included 205 IDHwt GBM patients, 24 cases of IDHmut grade 4 astrocytoma and 7 cases with unknown IDH status collected at Sahlgrenska University Hospital. All patients underwent standard of care treatment. All samples were collected during surgery and FFPE-embedded. Cohort 1 was used to investigate the presence of TLS in grade 4 glioma tissues, and to perform survival analysis based on the presence or absence of TLS in GBM patients. Inclusion criteria for the survival analysis were the following, (i) known IDH status; (iii) IDH status: wt; (iii) full survival information available, (iv) access to FFPE tumor blocks, leaving a total of 188 patients. The study was authorized by the Swedish Ethical Review Authority (DNR 2022-00160-01).

Surgically-resected samples used for all spatial transcriptomics assays were obtained under two different studies (Supplementary table 1, patients #237-242). Supratentorial glioblastoma samples from patients #237-239 (FFPE-embedded) and patients #241-242 (cryo-preserved) were identified in the database of the Department of Surgical Pathology, Uppsala University Hospital. The study was authorized by the regional Ethics Committee of Uppsala, Sweden (DNR 2015/089). The FFPE-embedded sample from patient #240 was collected during supramarginal glioblastoma resection at Sahlgrenska University Hospital and biobanked under a study approved by the Ethics Committee of Western Sweden (EPN/DNR: 559-12).

Surgically-resected samples used for single-cell sequencing (Supplementary table 1, patients #243-246) were obtained and collected during surgery at Uppsala University Hospital and Sahlgrenska University Hospital, under two studies approved by the regional Ethics Committee of Uppsala, Sweden (DNR 2007/353) and the Ethics Committee of Western Sweden (DNR 2022-00160-01).), respectively. All ethical regulations for work with human participants have been followed and informed consent was obtained. Analysis of tissues from all studies was authorized by the regional Ethics Committee of Uppsala, Sweden (DNR 2010/291).

### H&E assessment of patient samples

Tissues stained with routine hematoxylin-eosin (H&E) for pathological assessment at diagnosis were retrieved from the pathology departments of Sahlgrenska University Hospital or Uppsala University Hospital and scanned using high-throughput tissue scanners. Scanned tissues were independently assessed for the presence of lymphoid aggregates by at least one pathologist and one researcher with experience in tertiary lymphoid structure analysis. Visualization and manual annotation of H&E-stained tissue scans was performed using the ImageScope software (v 12.4.0.5043).

### Immunofluorescence staining of patient tissues

4µm-thick sections from FFPE-embedded samples were cut using a microtome and collected on Superfrost Plus Adhesion Microscope Slides (Epredia, #J1800AMNZ). Slides were incubated at 37°C overnight, baked at 60°C for 1 hour and deparaffinized using sequential incubations in Xylene (3 times x 10 minutes), 100% ethanol (1 time x 10 minutes), 95% ethanol (1 time x 10 minutes), followed by rinsing in 70% ethanol and distilled water (dH_2_O). Next, samples were incubated for 20 minutes in 10% neutral buffered formalin (NBF) and rinsed in dH_2_O. All deparaffinization steps were performed in slide chambers. Antigen retrieval was performed by boiling samples in AR6 buffer (Akoya Biosciences, #AR600250ML) in a heat-resistant jar using heating cycles of (i) 45 seconds at maximum power and (ii) 15 minutes at low power in a microwave. Slides were left for 15 minutes at room temperature, rinsed in dH_2_O and blocked using 1x Antibody diluent/Block solution (Akoya Biosciences, Opal 7-Color Manual IHC Kit, #NEL811001KT) for 30 minutes at room temperature. Slides were stained with primary antibodies diluted in 1x Antibody diluent/Block solution overnight at 4°C, washed with TBS-T (TBS + 0.05% Tween20) (3 times x 2 minutes) and incubated with secondary antibodies and Hoescht 33342 (Sigma-Aldrich, #14533) diluted in 1x Antibody diluent/Block solution for 2 hours at room temperature. Finally, slides were washed with TBS-T (3 times x 2 minutes), washed with dH_2_O (1 time x 2 minutes) and mounted using Fluoromount-G (Thermo Fisher Scientific, #00-4958-02).

10µm-thick sections from cryo-preserved OCT-embedded samples were cut using a cryostat and collected on Superfrost Plus Adhesion Microscope Slides (Epredia, #J1800AMNZ). Tissue was preserved at -80°C before use. Slides were thawed and dried at room temperature before fixation in acetone at 4°C for 10 minutes. Slides were removed from the acetone and let dry until all acetone had evaporated, followed by re-hydration in PBS for 5 minutes at room temperature and blocking with 1x Antibody diluent/Block solution (Akoya Biosciences, Opal 7-Color Manual IHC Kit, #NEL811001KT) for 1 hour at room temperature. Samples were incubated with primary antibodies diluted in 1x Antibody diluent/Block solution overnight at 4°C, washed with PBS (3 times x 5 minutes) and incubated with secondary antibodies and Hoescht 33342 (Sigma-Aldrich, #14533) diluted in 1x Antibody diluent/Block solution for 2 hours at room temperature. Finally, slides were washed with PBS (3 times x 5 minutes) and mounted using Fluoromount-G (Thermo Fisher Scientific, #00-4958-02).

The Vectra Polaris Automated Quantitative Pathology Imaging System (v1.0.13, Akoya Biosciences) was used to obtain 20x tile scans of the whole tissue slides.

Primary antibodies: mouse anti-human CD20 (clone L26, Dako); rabbit anti-human CD3 (clone SP7, Thermo Fisher Scientific); sheep anti-human CD34 (#AF7227-SP, R&D), rabbit anti-human coll IV (#ab6586, Abcam). Secondary antibodies: anti-mouse AF568 (Invitrogen, #A10037), anti-rabbit AF647 (Jackson ImmunoResearch, #711-605-152), anti-sheep 488 (Invitrogen, #A11015).

### Analysis of human tissues stained by immunofluorescence

Scans of human tissue samples stained with TLS markers (CD20, CD3, CD34 and nuclei) by immunofluorescence (IF) were manually screened to determine the presence of lymphoid aggregates (nr. of cells > 50). Tightly-packed lymphoid aggregates that contained T cells and/or B cells and exhibited a CD34^+^ vascular network were classified as TLS. Tightly-packed lymphoid aggregates that contained T cells and/or B cells and formed around a main vessel with a large lumen were classified as PNs. TLS and PNs were characterized as T cell-rich (T), mixed (M) or B cell-rich (B) based on the proportion of T cells and B cells within the aggregates. T-to-B cell proportions were visually assessed. The accuracy of the assessment was confirmed by comparing the results obtained by visual assessment with quantifications of the ratio of CD3^+^ area / CD20^+^ area within randomly-selected aggregates. B cell-rich aggregates that showed distinct T and B cell zones were classified as “zonated”. Nuclear density within TLS or PNs was quantified as nuclei area / total TLS area. Location of TLS and PNs was assessed using H&E-stained tissues to annotate whether each structure was proximal (<1500µm) or distal (>1500µm) from meningeal-rich areas of the brain, including the cortical surface, ventricular areas, gyri and sulci. Visualization of IF-stained slides and quantifications were performed using the QuPath software (version 0.4.3)^31^ and the ImageScope (version 12.4.0.5043).

### Cell culture

The murine glioma cell line CT-2A (developed by Thomas Seyfried, Boston College, Boston, MA) was cultured in Roswell Park Memorial Institute (RPMI) 1640 medium (#21875-034) supplemented with 10% (vol/vol) heat-inactivated fetal bovine serum (FBS) (#10082-147). All cell culture reagents were purchased from Gibco, Thermo Fisher Scientific. Cells were cultured at 37°C with 5% CO2 in a humidified cell incubator. Tests for mycoplasma contamination were performed routinely using MycoAlert Detection Kit (Lonza, Basel, Switzerland, #LT07-705).

### In-vivo experiments

6-8-week-old female C57BL/6 mice were used for orthotopic implantation of CT-2A glioma cells. Mice were anesthetized with 2% isoflurane and immobilized in a stereotaxic frame on a heated surface. A midline incision was made on the scalp and a hole was drilled in the skull at +1 mm anteroposterior and +1.5 mm mediolateral stereotactic coordinates from the bregma. 5x10^4^ CT-2A cells were intracranially (i.c) injected in 2μl DPBS (Thermo Fisher Scientific, #14190-144) with a Hamilton syringe at a depth of 2.7 mm. The incision was closed using Vetbond (3M, St. Paul, MN, #1469SB) and the mice were observed until full recovery from anesthesia on a heated surface. Mice were monitored daily and scored for tumor-related symptoms, such as hunched posture, lethargy, persistent recumbency and weight loss, according to the Uppsala University (Uppsala, Sweden) scoring system for animal welfare. Mice were sacrificed at pre-determined time points or at the humane endpoint (here referred to as “survival endpoint”), when the welfare score reached a maximum of 0.5. Mice were sacrificed via ketamine/xylazine overdose followed by intracardiac perfusion with 10mL of PBS (Thermo Fisher Scientific) and 10mL of 4% (wt/vol) paraformaldehyde (PFA) (Sigma-Aldrich, #158127). Brains were collected and processed for immunofluorescence analysis. For the CD4 depletion experiment, mice were intravenously treated with 200µg of rIgG2b (BioXCell, #BE0085) or αCD4 (BioXCell, #BE0003-1) antibodies two days before tumor implantation, and with 100µg of either antibody on days 4, 11 and 18 post-tumor implantation. Blood was collected on day 6 post-tumor implantation from the tail vein, to confirm CD4 T cell depletion by flow cytometry.

All mouse experiments were approved by the Uppsala County regional ethics committee (DNR 19429/2019), and were performed according to the guidelines for animal experimentation and welfare of Uppsala University. This study used C57BL/6NTac (Taconic M&B, Bomholt, Denmark) mice. All mouse studies were designed and performed according to either ARRIVE guidelines or accounting for the 3Rs principle. Sample size was determined based on previous experience with biological variation in the *in vivo* model used and accounting for the 3Rs principle.

### Flow cytometric analysis of murine blood samples

Blood was collected from the tail vein using capillary tubes (Sarstedt, #16.443.100). 25µl of blood was stained with CD4-FITC (Biolegend, #100510) and CD8-APC-Cy7 antibodies (BP Pharmigen, #561967), diluted 1:100 in FACS buffer (PBS + 0.5% BSA + 2mM EDTA), for 20 minutes at 4°C. Red blood cells (RBCs) were lysed using 1X RBC lysis buffer (eBioscience, #00-4300-54), cells were resuspended in FACS buffer and data was acquired on a CytoFLEX LX machine (Beckman Coulter).

### Immunofluorescence staining of murine samples

Brains collected after intracardiac perfusion were fixed overnight in 4% (wt/vol) PFA cryoprotected in 30% (wt/vol) sucrose (in PBS) for at least 48h, and cut into 80μm-thick sections using a vibratome. The cerebrum was virtually divided in eight segments from the forebrain to the midbrain, and tissue sections were collected on slides so that each slide would contain 8 representative parts of the entire cerebrum. Sections were then blocked in PBS containing 1% bovine serum albumin (BSA) and incubated with primary antibodies overnight at 4°C. Sections were washed in PBS and mounted using Fluoromount-G (Thermo Fisher Scientific, #00-4958-02). For imaging and analysis, fluorescence tile scans were captured with a Leica DMi8 microscope (Leica Microsystems) and confocal images were acquired in a scan format of 1024x1024 pixels with a Leica SP8 confocal microscope (Leica Microsystems). Primary antibodies used: rat anti-mouse B220-PE (clone RA3-6B2, Biolegend); rat anti-mouse CD3-BV241 (clone 17A2, BD Biosciences); rat anti-mouse CD45-AF647 (clone 30F-11, Biolegend); goat anti-mouse CD31 (clone AF3628, R&D Systems); donkey-anti-goat-AF647 (#A-21447, Thermo Fisher Scientific).

### Analysis of murine tissues stained by immunofluorescence

Tile scans of whole brain sections were visually assessed to identify the presence of TLS and PNs. TLS were defined as tight clusters of CD45^+^ immune cells containing CD3^+^ T cells and/or B220^+^ B cells forming around a CD31^+^ vascular network. PNs were defined as tight clusters of CD45^+^ immune cells containing CD3^+^ T cells and/or B220^+^ B cells forming around one main vessel with a large lumen. High resolution confocal images were used to quantify the T-to-B cell ratio within each TLS (CD3^+^ area / B220^+^ area) using the Fiji (v1.21) software. Each structure was allocated to a T cell-rich (T, ratio > 1.25), mixed (M, 0.85 < ratio < 1.25) or B cell-rich (B, ratio < 0.85) category accordingly.

### GeoMx spatial transcriptomics assay

FFPE-embedded GBM tissues (Supplementary table 1, patients #237-240) underwent the Human Whole-Transcriptome GeoMx assay pipeline (Nanostring), following the steps indicated by the manufacturer. Concisely, 4µm-thick FFPE tissue sections underwent incubation with UV-cleavable probes designed for comprehensive transcriptome coverage. Lymphoid aggregates or tumor areas were identified as distinct regions of interest (ROIs) through the use of morphology markers as guidance, such as CD3 (Origene, #UM000048BF), CD20 (Novus Biologicals, #NBP2-47840AF532) and CD31 (R&D Systems, #AF3628) antibodies as well as Syto13 (Thermo Fisher Scientific). Oligonucleotides specific to tissue-bound probes were UV-cleaved and isolated individually for each ROI, and subsequently subjected to sequencing on the Illumina platform for identification of the resolved mRNA-specific probes. Raw data was presented as counts per ROI.

Data was processed and analyzed using the Nanostring DSP software, R (v4.2.0) software, Partek® Flow® and/or Enrichr^32–34^. In brief, data was Q3-normalized and cell proportions within each ROI were deconvoluted using the SafeTME matrix^35^ and the SpatialDecon (v1.8.0) package in R. Q3-normalized data was used to run intra-sample differential expression analysis in Partek using the *Compute Biomarker* task, to identify genes that were upregulated in the tumor area versus the lymphoid aggregates, and to compile a list of tumor-upregulated genes per patient. Tumor-upregulated genes from each patient were merged into one list (*TumorUpGenes*). For inter-patient analysis, the RUV4 algorithm was run on the Q3-normalized data to correct for batch effects using the standR (v1.5.1) package in R. Genes included in the *TumorUpGenes* list were subtracted from the analysis, to remove patient-specific tumor cell-related background. Differential expression analysis was run in Partek using the *Compute Biomarker* task to identify genes that were specifically upregulated in T-PNs, T-TLS, M-TLS and B-TLS (threshold of p<0.01). Biomarkers of each group were inputted in Enrichr^32–34^ (GO_Biological_Processes_2023) to identify group-specific biological processes (p<0.05). Relative expression of group-specific biomarkers that appertained to relevant immune and cellular processes were visualized using a heatmap (Figure 2C).

### Padlock-probe targeted in situ sequencing

Cryo-preserved GBM tissues (Supplementary table 1, patients #241-242) underwent Padlock-probe targeted in situ RNA sequencing to visualize the expression of immune and stromal-related mRNAs, (Supplementary table 4). In brief, 10µm-thick cryo-preserved OCT-embedded tissue sections were incubated with mRNA-targeted Padlock probes as previously described^36,37^. Probes that hybridized to their targets were enzymatically ligated and amplified using rolling-circle amplification (RCA), and signal/coordinates of each rolling-circle amplification product (RCP) were resolved using fluorophore-bound detection probes across multiple detection cycles, according to the HybISS protocol^38^. Cell nuclei were stained with DAPI. Image processing was performed as described by Lee et al^36^, and visualization of mRNA location on the tissue was performed using the TissUUmaps software (v3.2.0.1)^39^.

### Single-cell sequencing of tumor-infiltrating T cells in surgically-resected GBM

#### Processing of fresh surgical samples

Tumor core samples from surgically-resected high-grade gliomas (Supplementary Table 1, patients #243-246) were collected from both Akademiska University Hospital (Uppsala, Sweden) and Sahlgrenska University Hospital (Gothenburg, Sweden). Samples from Akademiska University Hospital were collected in ice-cold Hybernate-A medium (Gibco, #A1247501) and processed immediately after surgery. Samples from Sahlgrenska University Hospital were collected in ice-cold MACS Tissue Storage Solution (Miltenyi Biotec, #130-100-008) and shipped overnight to Uppsala, where they were processed the following day. Samples were managed under a sterile laminar flow hood. Tissue was cut in small pieces using a scalpel, and was dissociated into a single cell suspension using the Human Tumor Dissociation kit (Miltenyi Biotec, #130-095-929) on a gentleMACS™ Octo Dissociator (Miltenyi Biotec) following the instructions provided by the manufacturer. Dissociated samples were diluted with 10 ml of MACS buffer (PBS plus 0.5% BSA) added with 2mM EDTA (Invitrogen, #AM9260G), 2mM L-Glut (Gibco, #25030081), 1mM sodium pyruvate (Gibco, #11360-070), 1X MEM NEAA (#Gibco, #11140-050), 1X MEM amino acids (Gibco, #11130-036). Samples were filtered through a 70 µm nylon strainer and centrifuged 300g x 5 min at 4°C to remove supernatant. Cell pellets were resuspended in 5 ml of 25% BSA in a 15ml Falcon tube, and centrifuged at 1350g (low brake =2) at 4°C to separate the myelin from the cell pellet. Myelin ring and supernatant were vacuumed away, and cell pellets were incubated in 2ml of 1X RCB lysis buffer (eBioscience, #00-4300-54) for 5 minutes at room temperature to lyse red blood cells. Samples were washed in PBS and viably frozen in FBS plus 10% DMSO overnight at -80°C. Samples were stored long-term at -150°C.

#### Single-cell library preparation and sequencing

mRNA whole transcriptome single-cell libraries of tumor-infiltrating immune cells were prepared using the BD Rhapsody™ platform (BD BioSciences). Myelin-depleted single-cell suspensions obtained from high grade glioma samples were sequentially labeled using the BD Human Sample Multiplexing Kit (Sample-tag) (BD Biosciences, #633781), the sorting antibody CD45-APC (#560973) and the viability stain Zombie Aqua (Biolegend, #423101) following the manufacturer’s protocol (BD Biosciences). The stained cell suspension was enriched for live CD45^+^ immune cells using a BD AriaIII flow sorter (BD Biosciences). Sorted live CD45^+^ cells from each patient were counted and pooled at equal ratios to achieve a total of approximately 20,000 cells in 620μL of ice-cold BD Sample Buffer (BD Biosciences, #664887). The pooled samples were loaded onto a BD Rhapsody cartridge (BD Biosciences, #633733) and mRNA from single cells was captured using Enhanced Cell Capture Beads according to the manufacturer’s protocol (BD Biosciences, #664887). cDNA from the captured mRNA was obtained by performing reverse transcription and treatment with Exonuclease I (BD Biosciences, #633773), and whole-transcriptome and Sample-tag libraries were obtained using the BD WTA amplification kit and protocol (BD Biosciences, #633801). The quality of the final libraries was assessed by using an Agilent 2200 TapeStation with HS D5000 ScreenTape (Agilent Technologies, Santa Clara, CA, #5067-5592) and concentrations were measured by a Qubit Fluorometer using the Qubit dsDNA HS kit (Thermo Fisher Scientific, #Q32854). This process was repeated a total of two times to obtain two libraries, each containing samples from two patients. The two final libraries were diluted to 2nM and pooled for paired-end sequencing 51+8+0+71 cycles in one NovaSeq 6000 S4 lane (Illumina, San Diego, CA) at the SNP&SEQ Technology Platform (Uppsala, Sweden).

#### Seven Bridges processing of single-cell sequencing data

Fastq files obtained from the SNP&SEQ Technology Platform were processed using the standard Rhapsody analysis pipeline (BD Biosciences) on Seven Bridges. In brief, reads R1 and R2 were filtered by dropping reads that were too short or had a base quality score of less than 20 to ensure high-quality reads. Reads R1 were used to identify cell label sequences and unique molecular identifiers, and reads R2 were mapped to the human reference genome release 29 (GRCh38.p12). All valid reads R1 and R2 were combined and annotated to the respective molecules and cells. Recursive substation error correction (RSEC) algorithm developed by the manufacturer (BD Biosciences) was used to correct for PCR and sequencing errors. Final expression matrices contained RSEC. Pooled samples were deconvoluted using Sample-tag reads. A cell was annotated as a singlet if the minimum read count for a given Sample-tag was attained and if more than 75% of the Sample-tag reads were derived from a single Sample-tag antibody. Multiplets were assigned if the count for two or more Sample-tag antibodies exceeded the minimum threshold. Cells were assigned as undetermined if they did not reach the criteria for either a singlet or multiplet. The final read depths for the two libraries were in the following range: mRNA 49391.18 to 56047.77 mean reads/cell and 92.63-92.81% sequencing saturation which were in the expected range of the manufacturer’s recommendations for targeted transcriptomics sequencing (BD Biosciences).

#### Single-cell sequencing analysis workflow

All downstream analyses of single-cell sequencing data were performed in Partek Flow build version 11.0.24.0414 (Partek Inc., Chesterfield, MI). For analysis, we used RSEC-adjusted molecule counts. Undetermined, multiplets, cells with >20% mitochondrial gene counts and cells with <1000 detected transcripts were filtered out from the analysis. Data were normalized as counts per 10,000, added an offset of 1 count, and Log2-transformed as recommended by the manufacturer (BD Biosciences), using the data normalization module in Partek Flow. Potential batch effects related to differences in sample collection were corrected using the Harmony algorithm (31740819). Dimensionality reduction using PCA followed by UMAP (default settings) was performed on all genes. Unsupervised clustering was performed using the Louvain clustering algorithm (using default settings, nearest neighbor type: K-NN, number of nearest neighbors: 30) and biomarkers for each cluster were computed using ANOVA tests (default settings) with Benjamini-Hochberg test to correct for false discovery rate (FDR) value ≤ 0.05 and log2 fold change ≥1.5. CD8 and CD4 T cell clusters, as well as subpopulation of CD4 T cells were annotated based on gene expression profiles (Supplementary Figure 6, Supplementary table 10). Finally, a Seurat package was used to compare each identified CD4 T cell subpopulation with all non-CD4 T cell clusters in our sc-seq dataset, to create a list of genes enriched in the CD4 T cell population (Sc-seq_CD4enriched_list) (Supplementary table 11).

### CosMx single cell spatial transcriptomics assay

#### Sample preparation and data collection

FFPE-embedded GBM tissues (Supplementary table 1, patients #237-240) underwent the Human 1000-plex CosMx mRNA assay pipeline (Nanostring), following the steps indicated by the manufacturer (Supplementary table 5). Concisely, 4µm-thick FFPE tissue sections underwent incubation with antibodies and dyes specific to morphological markers essential for segmentation, such as 18S-RNA (Nanostring, #121500124) to resolve cell cytoplasm, and H3 (Nanostring, #121500125) and DAPI (Nanostring, #121303304) to resolve the nuclei. Fields of view (FOVs) were selected based on staining for CD3, CD20 and CD34 performed on a sequential section, with antibodies indicated under the “Immunofluorescence staining of patient tissues” paragraph of the Methods section of this paper. Probe signals/coordinates were acquired through successive cycles of fluorescence *in situ* hybridization (FISH), and attributed to individual cells based on cell segmentation. Raw data was presented as counts per cells.

#### Data QC, normalization and annotation of cell clusters

Data was processed and analyzed using the R (v4.2.0) software. Quality control (QC) was performed individually for each sample. Only cells that fulfilled the following requirements and passed QC were included in the downstream analysis (Supplementary Table 12): (i) counts of negative probes/cell < 0.5 (to remove cells with high background signal); (ii) probe counts/cell < 2000 and (iii) cell size < 5 times the standard deviation of the average cell size (to remove potential doublets); (iv) nr of detected mRNAs/cell > 20 (to remove low quality cells). Data from all samples was normalized using SCtransform and pre-processed using a principal component (PCA) analysis to remove noise. Cell clusters were identified using the Louvain algorithm and visualized on a UMAP. Batch effects were removed using the Harmony algorithm, and the PCA → Louvain clustering → UMAP steps were repeated to identify the batch-corrected cell clusters (Supplementary Figure 4A). Annotation of cell clusters was done using gene expression scores based on the gene signatures in Supplementary Table 6 (Supplementary Figure 4B-J), and by manually checking that the most expressed genes in each annotated cluster corresponded to their assigned cell type. Cluster 4 (Supplementary Figure 4A) contained marker genes of both macrophages and fibroblasts, thus clusters 2, 4, 9 and 13 were extracted and re-clustered (following the above listed steps) to improve the output of cell type identification, obtaining the final annotations shown in Figure 4J. The SingleCellExperiment (v1.20.1) package for R was used for step (i) of the QC, while the Seurat (v4.3.0) package for R was used for steps (ii-iv) of the QC, as well as for the PCA, Louvain clustering and UMAP visualization. The Harmony (v1.0.3) package for R was used for batch effect removal. The Ucell (v2.2.0) package for R was used to plot the gene expression scores.

#### Identification of CD4 and CD8 T cell clusters in the CosMx dataset by label transfer

scRNA-seq data obtained for the T cell population was used as reference to perform the label transfer analysis on the CosMx dataset. In brief, after annotating CD4 T cells and CD8 T cells in our scRNA-seq cohort, we calculated the average expression of the 990 genes included in the CosMx gene panel within these populations. The Insitutype package (v 1.0.0) in R was then used to do the label transfer analysis^40^. In brief, the average gene expression profile of CD4 T cells and CD8 T cells from the scRNA-seq dataset was loaded in to R, together with the expression count matrix of the T cell population from the CosMx dataset (as well as the corresponding mean negative control value of each T cell). Next, we applied a supervised cell typing model from the Insitutype package called “insitutypeML” to achieve the following analysis steps: (1) adjust the expression value of each T cell by using the average negative control value and (2) calculate the loglikelihood of the adjusted count matrix of T cells from the CosMx dataset, using our reference scRNA-seq data to assign the T cell subtypes automatically (CD4 or CD8). *Spatially-resolved cell-cell interaction analysis*

The SpaTalk package (v1.0) in R was used to perform the cell-cell interaction analysis^41^. Ligand– receptor interactions (LRIs) from CellTalkDB, pathways from KEGG and Reactome, and transcription factors (TFs) from AnimalTFDB were integrated to construct the ligand–receptor–TF knowledge graph (LRT-KG) in this package. SpaTalk objects including the expression matrix, spatial coordinates and cell type annotation were created for each investigated FOV. Next, the Euclidean distance between cells in each FOV was calculated using the single-cell spatial coordinates, and the KNN algorithm was used to select the 10 nearest neighbors of each cell to construct the cell graph network and set the “cell-cell pairs”. The number of cell-cell pairs expressing ligand and receptor was obtained using the graph network, after which an inter-cellular score and p-value (threshold p<0.05) were calculated for each LRI to identify non-random predicted interactions within each FOV, using a permutation test to filter and score the significantly enriched LRIs. Finally, the random-walk algorithm was applied to filter and score the expression of TFs and targets downstream of each significant predicted interaction.

#### Trajectory analysis of CD4 T cells

To compute a list of gene relevant to interrogate CD4 T cell states in GBM tissues, we exploited single-cell sequencing data of tumor-infiltrating immune cells isolated from surgically-resected high grade gliomas (see the paragraph “Single-cell sequencing of tumor-infiltrating T cells in GBM”). A list of genes that was highly expressed in CD4 T cells compared to other tumor-infiltrating immune cells in our sc-seq dataset (Sc-seq_CD4enriched_list) was created as indicated in the paragraph “Single-cell sequencing of tumor-infiltrating T cells in GBM”, subparagraph “Single-cell sequencing analysis workflow”. Next, the aforementioned list was compared with the CosMx gene list (Supplementary table 5) to identify overlapping genes that could be interrogated in our CosMx cohort. Lastly, manual curation of the list was performed, to include genes that were necessary to discriminate T helper states or relevant for lymphoid development, obtaining a final “CD4_trajectory_list” of 137 genes (Supplementary table 9). Trajectory analysis of CD4 T cells was then performed inputting the “CD4 trajectory list” and using the Monocle (v2.26.0) package in R. First, all CD4 T cells included in FOVs annotated as “Tumor”, “TLS” or “PN” in the CosMx dataset were extracted and assigned into five different groups, depending on their region of origin in the tissue (Tumor, T-PN, T-TLS, M-TLS, B-TLS). Next, the expression level of the marker genes included in the “CD4 trajectory list” were used as reference to infer the trajectory of CD4 T cells in all regions by using a Monocle standard pipeline, obtaining six main CD4 T cell states (Figure 7D,E). The ‘residualModelFormulaStr =c(“∼Sample”)’ parameter was set to remove the batch effect among different samples when performing the reduceDimension step. Finally, hierarchical clustering of genes was visualized with a heatmap (Supplementary Figure 7), and the different CD4 states were annotated based on gene expression.

### Statistical analysis

Statistical analysis was performed with GraphPad (v9.0.0), Partek® Flow® or R (v4.2.0). In GraphPad, Kaplan-Meier survival curves were analyzed using the Log-Rank test. Normal distribution of the data was investigated using either a D’Agostino and Pearson normality test or a Shapiro-Wilk normality test. When comparing two groups, a parametric *t*-test was used for normally distributed data, while a Mann-Whitney test was used for non-normally distributed data. When comparing three or more groups, a one-way ANOVA (correction for multiple comparisons: Tukey’s) was employed if the data was normally distributed, while a Kruskal-Wallis test (correction for multiple comparisons: Dunn’s) was used for non-normally distributed data. Statistics in Partek® Flow® and R workflows was performed according to the statistical tests within each task or package.

## Acknowledgments

This work was funded by the Swedish Cancer Society (20 1008 PjF, 20 1010 UsF, 23 3098 Pj), the Swedish Childhood Cancer Society (PR2021-0122), the Swedish Research Council (2020-02563), the Knut and Alice Wallenberg Foundation (KAW 2019.0088), Hjärnfonden (FO2022-0366) and the Swedish state (ALF-agreement, ALFGBG-965622). ICF was funded by Anillo InflamAIDS (ATE220016) ANID, Chile.

## Author contributions

AV and ADi conceptualized the project; AV, TvdW, SF, RL, ADe, AM, ICF and MR performed experiments; AV, FY, SF and MR performed data analysis; AV and FY curated and visualized the data; AS, SL, LU, FL, TOB and ASJ provided human material and clinical data; AV, ADe, SL and TOB assessed surgical tissues; ME, MN and LH provided input and resources for advanced methodology and analysis; AV wrote the original draft and produced the figures; All authors reviewed the manuscript; ADi provided funding and supervised the project.

