## Supplemental information for "Cellular and molecular events organizing the assembly of tertiary lymphoid structures in glioblastoma"

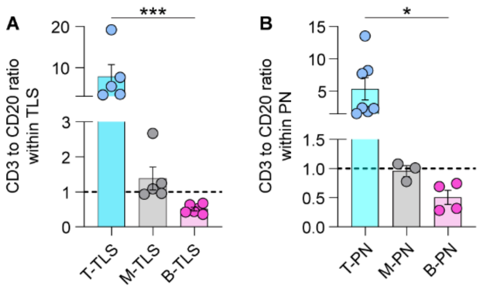


**Supplementary Figure 1. Characterization of the lymphocytic composition of TLS and PNs in human grade 4 astrocytoma tissues (including GBM and IDHmut grade 4 astrocytoma tissues).** ***Related to Figure 1****.* **(A-B)** CD3-to-CD20 ratio (CD3+ area / CD20+ area) was calculated within randomly-selected (A) TLS and (B) PNs, after visual classification of these aggregates as T cell-rich (T), mixed (M) or B cell-rich (B). The graph shows good correlation between the visual classification (x axis) and the calculated CD3/CD20 ratio (y axis). n=3-7 aggregates/group. Bar graphs show mean ±SEM. Statistics: (A) Kruskal-Wallis test, (B) one-way ANOVA. *p<0.05; ***p<0.001.

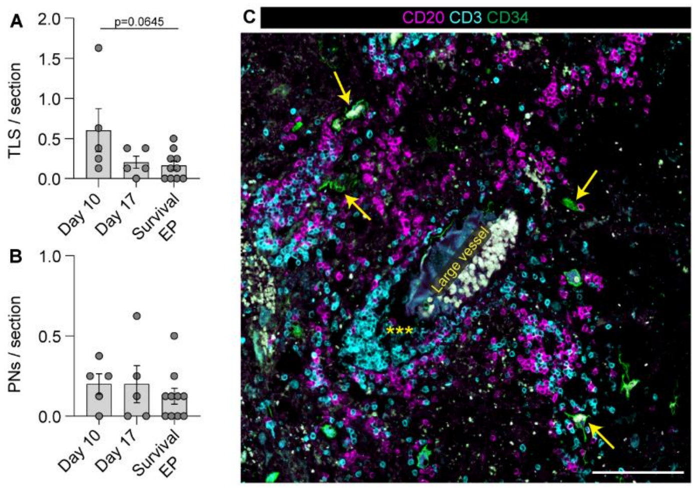


**Supplementary Figure 2. Characterization of TLS and PNs in murine and human GBM.** ***Related to Figure 3.*** **(A)** Number of TLS per section at the indicated time points post-tumor implantation. n=5-10 mice/group; Statistics: one-way ANOVA with Tukey's correction for multiple comparisons. **(B)** Number of PNs per section at the indicated time points post-tumor implantation. n=5-10 mice/group; Statistics: Kruskal-Wallis test with Dunn's correction for multiple comparisons. Bar graphs in (A,B) indicate mean ±SEM. **(C)** Representative immunofluorescence image of a large vessel surrounded by a T-PN in human GBM tissue (*** indicates T cell aggregation within the T-PN). Additional areas of dense lymphocyte clustering can be observed in the vicinity, which were associated with smaller vasculature (yellow arrows). Scale bar: 100µm.

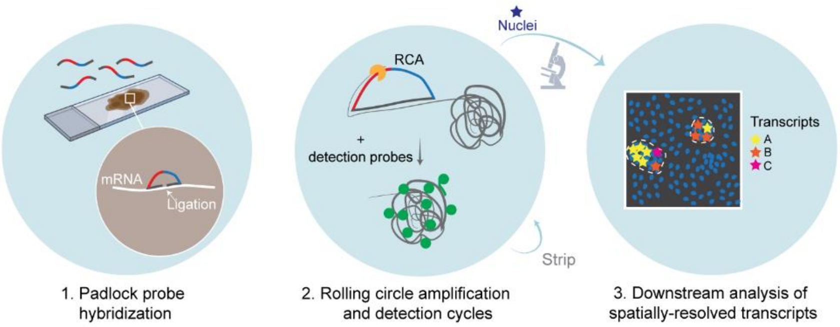


**Supplementary Figure 3. Schematic representation of the main steps performed for targeted Padlock probe-based *in situ* sequencing. *Related to Figure 4*. (A)** Schematic representation of the layout used for the targeted Padlock probe-based *in situ* sequencing assay. In brief, cryosections of human GBM tissue were stained with targeted Padlock probes. Probes that bound to their targets were amplified using enzymatic rolling circle amplification (RCA) and visualized using fluorescent probes. Probe signal was detected using cycles of fluorescence imaging and probe location on the tissue was recorded using coordinates. Cell nuclei were visualized using a nuclear dye.

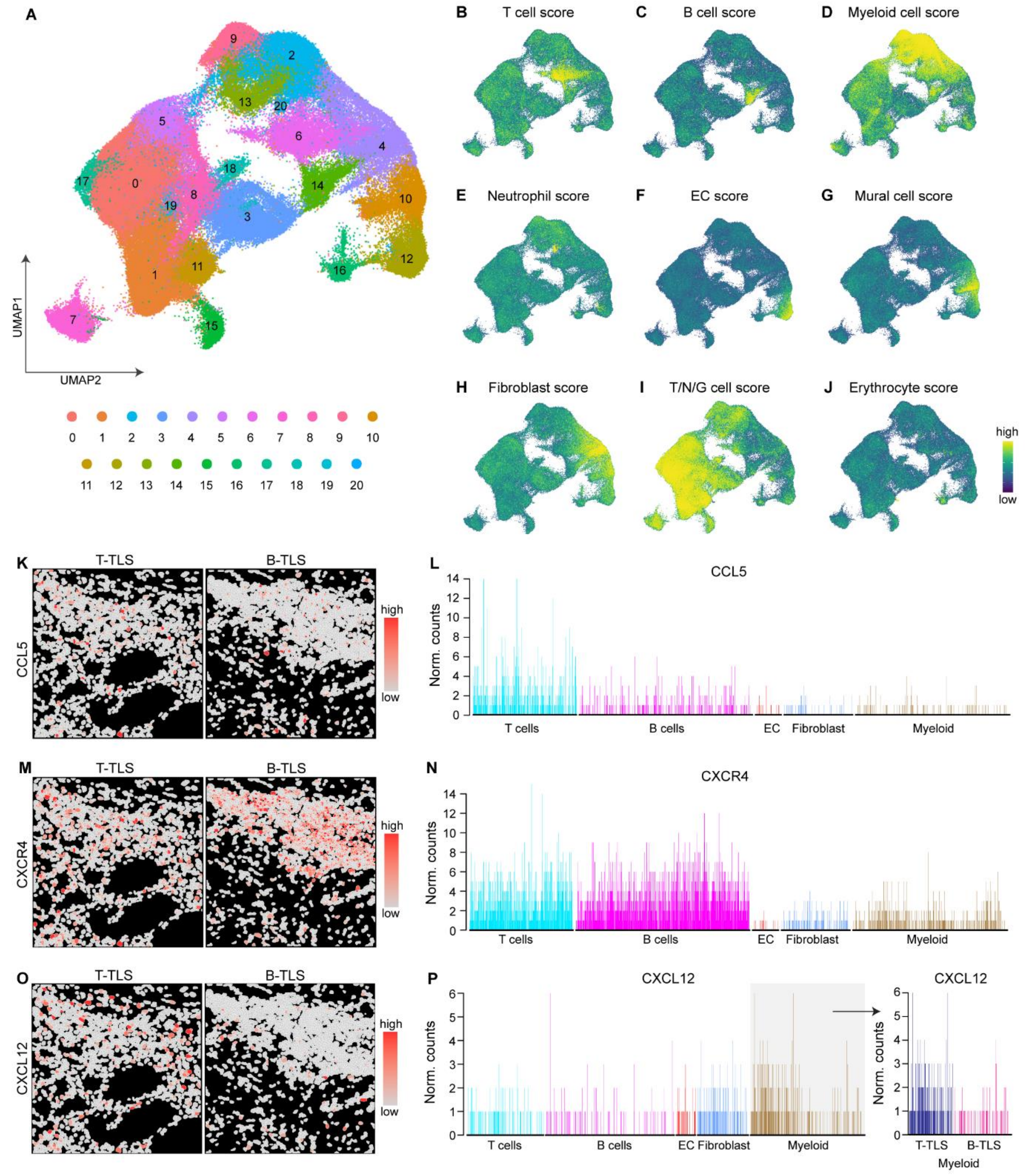


**Supplementary Figure 4. Characterization of cell types and gene expression within TLS using CosMx single-cell spatial analysis.** ***Related to Figure 4*****. (A)** UMAP of cell clusters identified in GBM tissues using the CosMx technology. n=4 patients. **(B-J)** Visualization of the cell scores used to annotate the indicated cell types onto the UMAP: (B) T cell score; (C) B cell score; (D) Myeloid cell score; (E) Neutrophil score; (F) Endothelial cell (EC) score; (G) Mural cell score; (H) Fibroblast score; (I) Tumor/Neuron/Glia (T/N/G) cell score; (J) Erythrocyte score. **(K,L)** Expression of CCL5 in single cells within T-TLS and B-TLS visualized as (K) an expression heatmap or (L) a bar graph displaying normalized counts of CCL5 in each cell within the shown FOVs. **(M,N)** Expression of CXCR4 in single cells within T-TLS and B-TLS visualized as (M) an expression heatmap or (N) a bar graph displaying normalized counts of CXCR4 in each cell within the shown FOVs. **(O,P)** Expression of CXCL12 in single cells within T-TLS and B-TLS visualized as (O) an expression heatmap or (P) a bar graph displaying normalized counts of CXCL12 in each cell within the shown FOVs. Grey box indicates bimodal distribution of CXCL12 expression in the myeloid cell population, related to differences in T-TLS vs B-TLS.

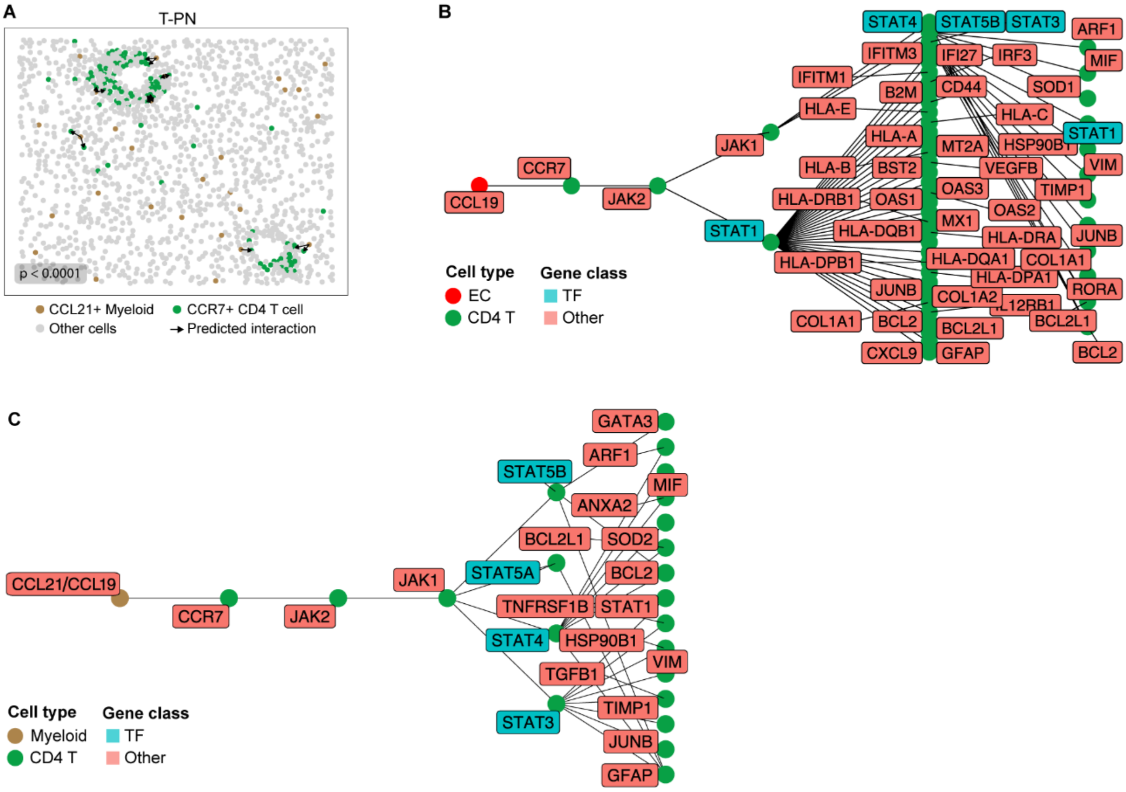


**Supplementary Figure 5. Ligand-receptor interactions between CD4 T cells and stromal/immune cells in T cell-rich aggregates in human GBM.** ***Related to Figure 6*****. (A)** Interaction plot showing significant predicted interactions between CCR7$^{+}$ CD4 T cells and CCL21$^{+}$ myeloid cells in T-PNs. Distribution of major cell types in the FOV shown in this panel can be found in Figure 5C. **(B)** Pathway-related results of the interaction analysis displayed in panel (A), showing that CCR7$^{+}$ CD4 T cells that were predicted to interact with CCL19$^{+}$ ECs express genes located downstream of the CCL19-CCR7 signaling cascade. **(C)** Pathway-related results of the interaction analysis displayed in Figure 6E,F, showing that CCR7$^{+}$ CD4 T cells that were predicted to interact with CCL19$^{+}$ or CCL21$^{+}$ myeloid cells express genes located downstream of the CCL19/CCL21-CCR7 signaling cascade. In (B,C), TF= transcription factor.

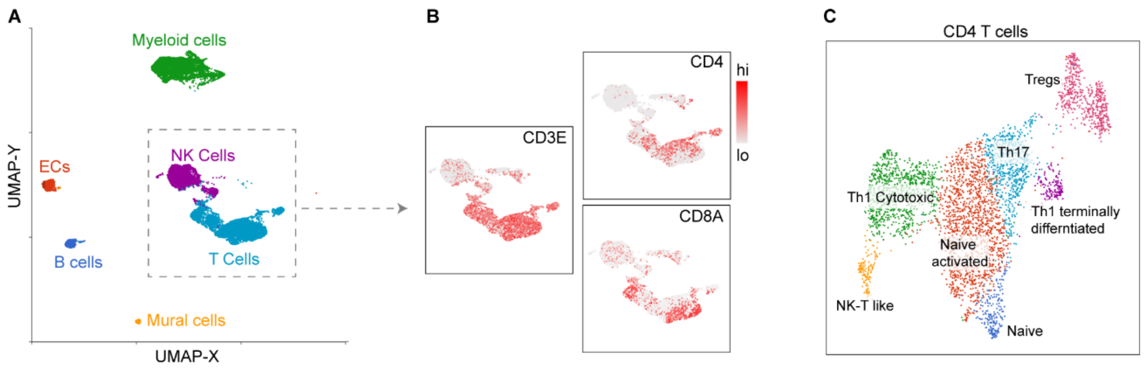


**Supplementary Figure 6. Identification of GBM-infiltrating CD4 T cells in surgically-resected GBM tissues via single-cell sequencing.** ***Related to Figure 7.*** **(A)** Major cell clusters identified via single-cell sequencing of GBM tissues, in immune cell-enriched samples. **(B)** Expression of CD3E, CD4 and CD8A within the T cell cluster. **(C)** Annotation of CD4 T cell subpopulations identified in this cohort (for cluster-specific gene expression, see Supplementary Table 10).

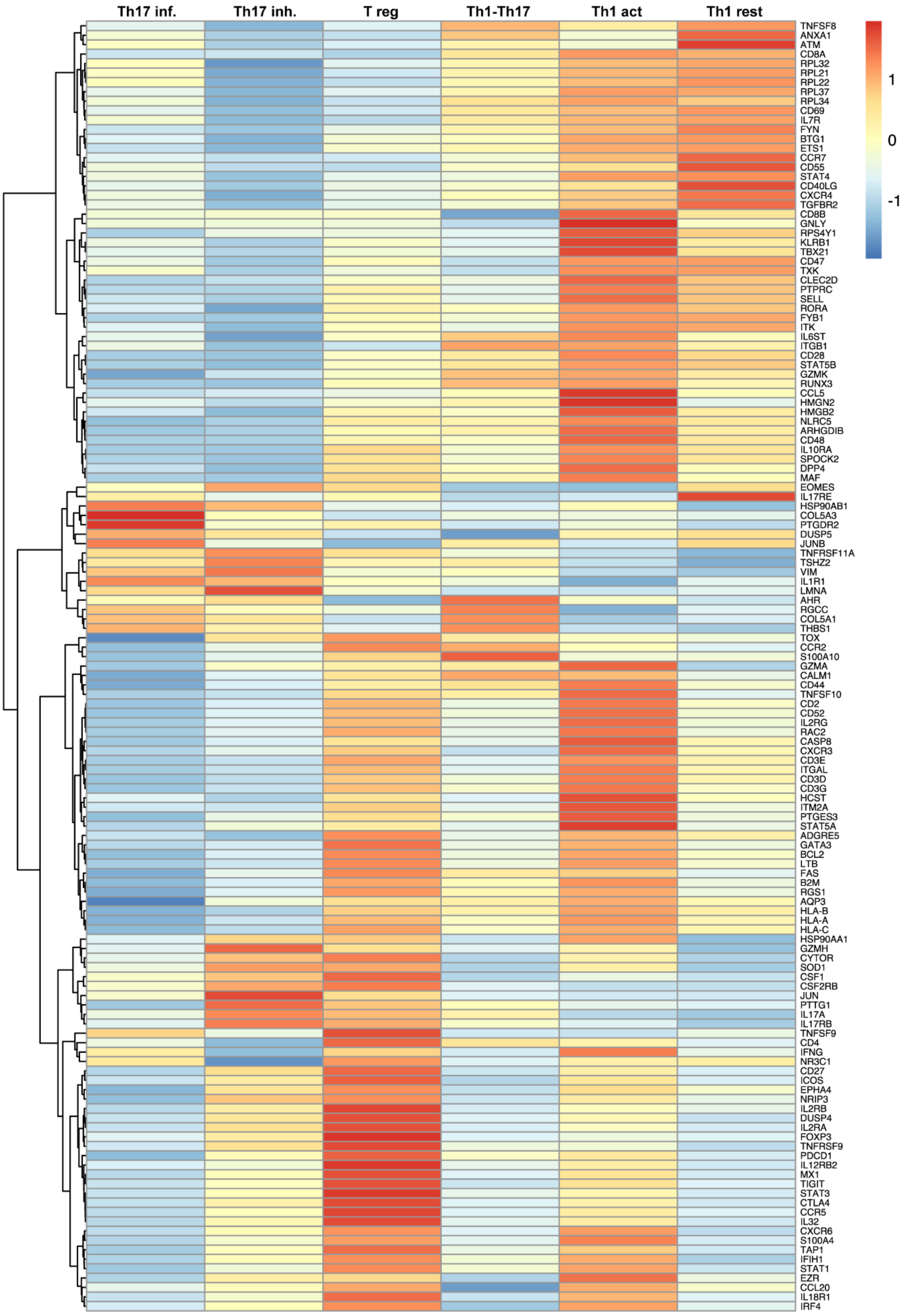


**Supplementary Figure 7. Relative gene expression of CD4 T cell subpopulations infiltrating GBM tumors.** ***Related to Figure 7*****.** Heatmap displaying the relative expression levels of markers included in the CD4_Trajectory_list (Supplementary table 9) across the six CD4 T cell subpopulation identified in Figure 7E (CosMx dataset). Th17inf = Th17 inflammatory; Th17inh= Th17 inhibitory; Treg = T regulatory cell; Th1act = Th1 activated; Th1 rest = Th1 resting.

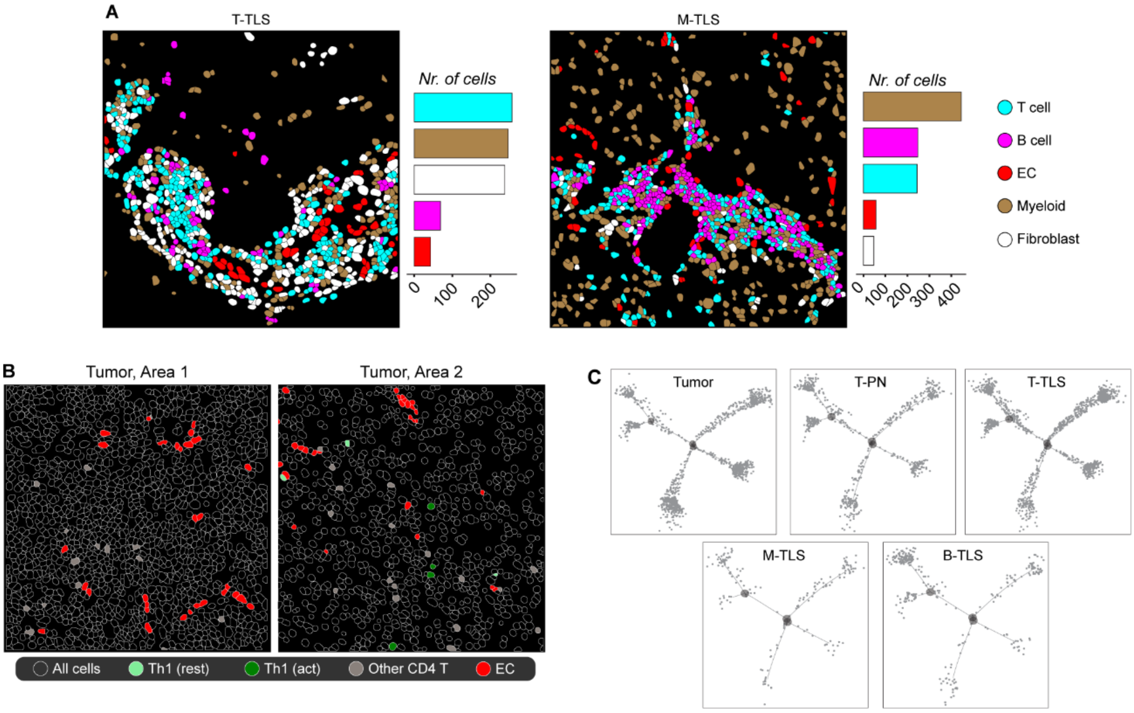


**Supplementary Figure 8. Characterization of T cells infiltrating lymphoid aggregates or tumor in human GBM using the CosMx single-cell spatial transcriptomics assay. *Related to Figure 7*. (A)** Spatial organization of T cells, B cells, endothelial cells (ECs), myeloid cells and fibroblasts in the T-TLS and M-TLS FOVs shown in Figure 7H. **(B)** Representative images of the spatial distribution of the Th1(rest) and (Th1act) CD4 T cell subpopulations in the tumor. **(C)** Trajectory plots showing the distribution of CD4 T cells across the six CD4 states identified in Figure 7E, in the indicated areas of the tissue (tumor, T-PNs, T-TLS, M-TLS and B-TLS).

### Supplementary tables - Legends

**Supplementary table 1.** Information about patients included in the current study, such as diagnosis, survival, IDH status, TLS status, sample type and assay performed. The dash symbol (-) indicates “no”, the plus (+) symbol indicates “yes”.

**Supplementary table 2**. ***Related to Figure 2.*** *Related dataset: GeoMx.* Full list of biomarker genes of T-PNs, T-TLS, M-TLS, B-TLS and B-PNs identified by GeoMx spatial transcriptomics analysis of human GBM tissues from patients #237-240.

**Supplementary table 3**. ***Related to Figure 2.*** *Related dataset: GeoMx.* Most relevant group-specific biological processes found to be enriched in T-PNs, T-TLS, M-TLS and B-TLS in the GeoMx dataset (patients #237-240) using Enrichr analysis.

**Supplementary table 4**. ***Related to Figure 4.*** List of mRNA probes used for Padlock probe-based *in situ* sequencing of human GBM tissues from patients #241-242.

**Supplementary table 5**. ***Related to Figure 4-7.*** *Related dataset: CosMx.* List of mRNA targets included in the CosMx spatial transcriptomics panel used in this study.

**Supplementary table 6**. ***Related to Figure 4.*** *Related dataset: CosMx.* Gene signatures used to annotate major cell clusters obtained via CosMx spatial analysis of human GBM tissues from patients #237-240.

**Supplementary table 7**. ***Related to Figure 6.*** *Related dataset: CosMx.* Differentially expressed genes in CD4 T cells located in spatially-resolved areas of GBM tissue from patient #240 (PNs, TLS or tumor).

**Supplementary table 8**. ***Related to Figure 6-7.*** *Related dataset: CosMx.* Differentially expressed genes in CD4 T cells vs CD8 T cells infiltrating GBM tissue from patients #237-240.

**Supplementary table 9**. ***Related to Figure 7.*** *Related dataset: CosMx.* List of genes used to perform the trajectory analysis of CD4 T cells (CD4_trajectory_list).

**Supplementary table 10**. ***Related to Supplementary Figure 6.*** *Related dataset: Sc-seq.* Biomarkers of CD4 subpopulations identified by single-cell sequencing in GBM tissues from patients #243-246.

**Supplementary table 11**. ***Related to Supplementary Figure 6.*** *Related dataset: Sc-seq.* List of genes enriched in of CD4 T cells versus all other tumor-infiltrating immune cells clusters identified in our single-cell sequencing dataset (patients #243-246).

**Supplementary table 12**. ***Related to Figure 4-7.*** *Related dataset: CosMx.* Sample-wise quality control (QC) parameters of GBM samples included in the CosMx spatial assay (patients #237-240).
